# Exuberant *de novo* dendritic spine growth in mature neurons

**DOI:** 10.1101/2023.07.21.550095

**Authors:** Sarah Krüssel, Ishana Deb, Seungkyu Son, Gabrielle Ewall, Minhyeok Chang, Hey-Kyoung Lee, Won do Heo, Hyung-Bae Kwon

## Abstract

Dendritic spines are structural correlates of excitatory synapses maintaining stable synaptic communications. However, this strong spine-synapse relationship was mainly characterized in excitatory pyramidal neurons (PyNs), raising a possibility that inferring synaptic density from dendritic spine number may not be universally applied to all neuronal types. Here we found that the ectopic expression of H-Ras increased dendritic spine numbers regardless of cortical cell types such as layer 2/3 pyramidal neurons (PyNs), parvalbumin (PV)- and vasoactive intestinal peptide (VIP)-positive interneurons (INs) in the primary motor cortex (M1). The probability of detecting dendritic spines was positively correlated with the magnitude of H-Ras activity, suggesting elevated local H-Ras activity is involved in the process of dendritic spine formation. H-Ras overexpression caused high spine turnover rate via adding more spines rather than eliminating them. Two-photon photolysis of glutamate triggered *de novo* dendritic spine formation in mature neurons, suggesting H-Ras induced spine formation is not restricted to the early development. In PyNs and PV-INs, but not VIP-INs, we observed a shift in average spine neck length towards longer filopodia-like phenotypes. The portion of dendritic spines lacking key excitatory synaptic proteins were significantly increased in H-Ras transfected neurons, suggesting that these increased spines have other distinct functions. High spine density caused by H-Ras did not result in change in the frequency or the amplitude of miniature excitatory postsynaptic currents (mEPSCs). Thus, our results propose that dendritic spines possess more multifaceted functions beyond the morphological proxy of excitatory synapse.

## INTRODUCTION

Dendritic spines are small membrane protrusions stemming from the dendrite of neurons. They serve as a structural compartment for the efficient information transfer from one neuron to another. During learning or normal development, these synapses undergo reorganization to establish functionally specific neural circuits ^1–9^. Since most excitatory synapses are built on these dendritic spines in mammalian brain, examining dendritic spine dynamics has been regarded as an anatomical proxy for changes in neural connectivity. Because of the strong coupling between dendritic spines and functional synapses, many molecules involved in spine number changes were identified as cell adhesion molecules ^10–13^.

Evidence of producing dendritic spines without having potential involvement of presynaptic neurons was also reported. Focal, repetitive release of glutamate near the dendrite by two-photon photolysis was able to generate *de novo* dendritic spines in layer 2/3 PyNs of the somatosensory cortex ^14, 15^. This glutamate-induced spinogenesis was not just limited to the somatosensory cortex but also found in other cortical brain areas such as motor cortex and prefrontal cortex ^16, 17^. Hippocampal PyNs and medium spiny neurons in the striatum also showed a similar spine forming process ^18, 19^. These data indicated that glutamate itself is a sufficient trigger for spinogenesis. However, when glutamate was photoreleased, spine formation was successful in one dendritic region but failed in another region even if glutamatergic receptor density was identical within the same neuron, even on the same dendritic branch ^14^. This observation suggests that what determines successful spinogenesis may not be the amount of glutamate nor the postsynaptic receptor density. Furthermore, γ-aminobutyric acid (GABA), an inhibitory neurotransmitter, also produced dendritic spines ^15^, indicating that the determining factor for spine formation is not the identity of neurotransmitter but presents in the postsynaptic side.

Another line of evidence supported that dendritic spine formation is independent of presynaptic glutamate or GABA release. Dendritic spines were normally formed and maintained in a mouse mutant lacking Munc13-1/Munc13-2 (M13 double knockout [M13 DKO]) where both glutamate and GABA release are completely abolished ^20^. These data suggest that spine formation occurs independent of presynaptic activity, and individual cells have a molecular machinery that can generate dendritic spines in a cell-autonomous manner. Another genetic manipulation that abolished vesicular glutamate release also showed normal spine density ^21^. Whether these activity-independently formed spines are able to make functional synapses is still unknown but their morphology was not different from functional spines and their ultrastructure showed normal contact to the presynaptic boutons ^20–22^. Evidence that glutamatergic neurotransmission is unnecessary for dendritic spine formation was also shown by selective deletion of ionotropic glutamatergic receptors in hippocampal CA1 PyNs ^23^. Thus, it seems apparent that dendritic spines are induced in the absence of neurotransmitters or receptor activation.

Despite these findings, what remains unclear is which molecules directly mediate dendritic spine protrusions and if so, whether these spines always become functional synapses. In adulthood, learning induces new dendritic spines not randomly but clustered with new spines being added in close proximity to preexisting stable spines involved in the task ^1, 24–28^. Preexisting spines exhibit signs of recent structural plasticity suggesting that signaling molecules spread from the stimulated spine to nearby dendritic segments priming the area for *de novo* spine formation^14, 24, 25^. One candidate as spreading, spinogenic molecule is the small GTPase H-Ras, which has been identified to be activated and stay activated after long-term potentiation (LTP) for up to 5 minutes and travels up to ∼10 μm away from the induction site ^24, 29, 30^. Additionally, in an earlier study success of glutamate induced *de novo* spine formation was reduced when H-Ras downstream signaling, mitogen-activated kinase (MAPK) kinase 1/2 (MEK1/2), was inhibited ^14^.

Here, we found that H-Ras overexpression increased dendritic spine number in excitatory PyNs as well as two different types of INs, PV-INs and VIP-INs, which normally do not express high density of dendritic spines ^31–39^. When investigating spine dynamics, H-Ras did not affect spine elimination rates but increased the probability of new dendritic spine formation in the presence and absence of external activity. These findings imply that H-Ras functions as a spinogenic molecule that produces dendritic spines independent of cell types. We uncovered that these exuberant number of spines are not coupled with functional connectivity changes. These data suggest that the function of dendritic spines need to be understood in a broader sense.

## RESULTS

### H-Ras increases dendritic spine number in pyramidal cells

We investigated the role of H-Ras on spinogenesis of mature PyNs in layer 2/3 of the neocortex. Recently developed intensiometric small GTPase biosensors allow for simultaneous overexpression and visualization of small GTPase activity patterns ^40^. In this study, we used the intensiometric small GTPase biosensor for H-Ras overexpression and activity detection. In its active state H-Ras interacts with its effector domain RAF initiating downstream signaling ^41^. This feature of heterodimerization is here coupled with the heterodimerization of two quenched fluorescent protein-derived monomers of ddFP, copy A and copy B, which produce bright green fluorescence when reunited (Figure 1A). Copy A was bound to H-Ras and copy B to Ras-binding domain (RBD_RAF_) sensing H-Ras activity upon H-Ras/ RBD_RAF_ binding. As controls, we created both a biosensor version devoid of its H-Ras component (from here on referred to as “RBD_RAF_”) and a biosensor version containing only the biosensor backbone without any signaling proteins (from here on referred to as “ΔH-Ras”) (Figure 1A).

**Figure 1:**
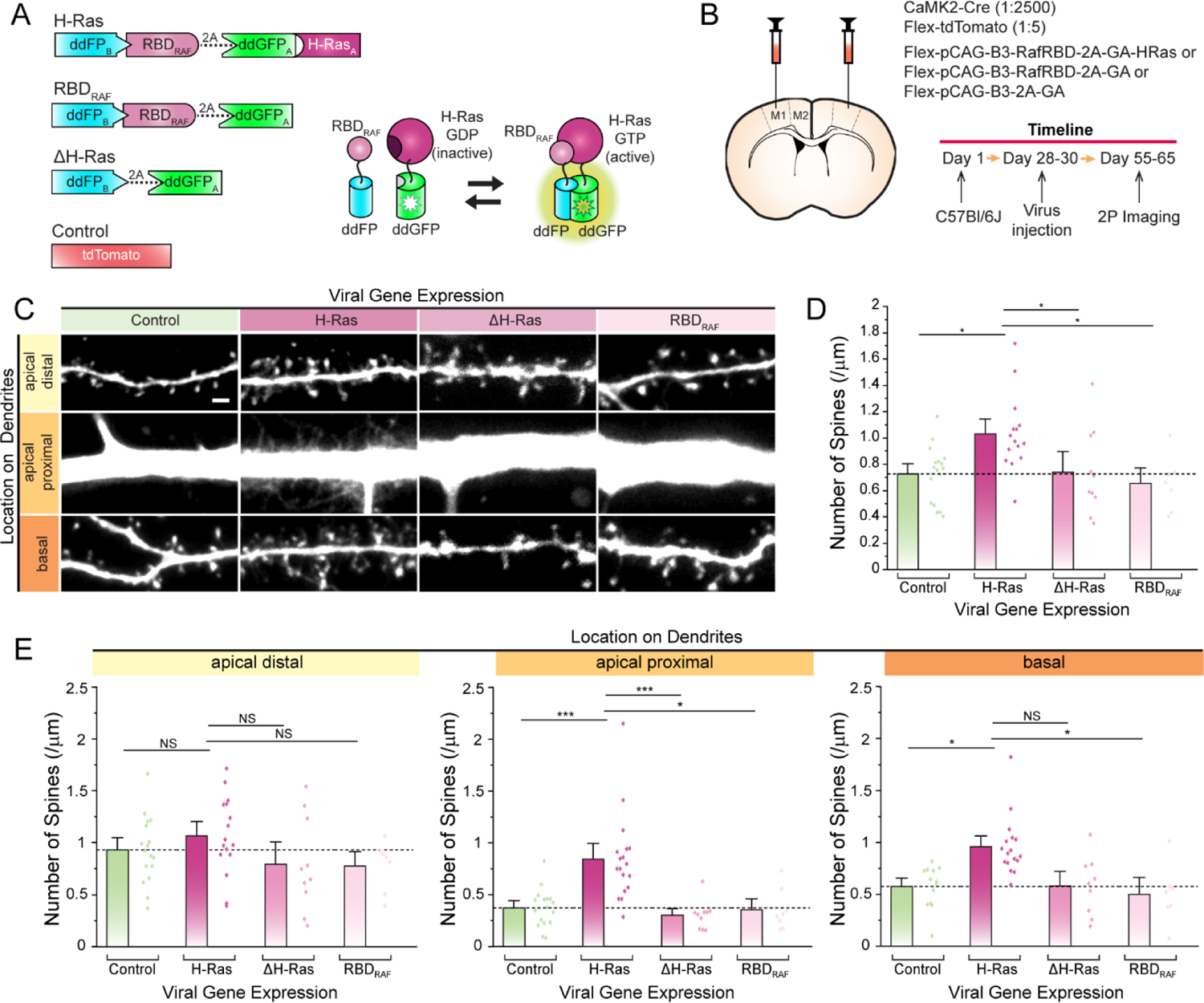
H-Ras increases spine number in cortical pyramidal neurons. **(A)** Left: Schematic depiction of H-Ras sensor constructs and its controls (RBD_RAF_, ΔH-Ras, and Control). Right: Schematic of the mode of action of ddFP-based H-Ras sensor. **(B)** Virus injection scheme and experimental timeline. **(C)** Two-photon microscopy images showing representative dendrites (apical distal, apical proximal, and basal) of pyramidal neurons expressing Flex-tdTomato in the primary motor cortex of a typical acute brain slice made from C57Bl6 mice at ∼P60 injected 4-weeks prior with CaMKII-Cre, Flex-tdTomato and either Flex-pCAG-B3-RafRBD-2A-GA-HRas (H-Ras), Flex-pCAG-B3-RafRBD-2A-GA (RBD_RAF_) or Flex-pCAG-B3-2A-GA (ΔH-Ras). Scale bar, 2 μm. **(D)** A summary graph showing the spine density. Dots represent average spine number from each neuron (dots) and bars indicate mean ± SEM, respectively for each condition (Control: 0.7259 ± 0.05162, N=17; H-Ras:1.0316 ± 0.0743, N=15; ΔH-Ras: 0.73881 ± 0.10413, N=10; and RBD_RAF_: 0.6554 ± 0.07848, N=7). The dotted line represents the average spine density of the tdTomato only control. *p<0.05 (Two-way ANOVA) **(E)** Separation of spine density analysis into the three dendritic locations: apical distal, apical proximal and basal. Apical distal: Control (0.931 ± 0.0776), H-Ras (1.06549 ± 0.09451), ΔH-Ras (0.79241 ± 0.14282), and RBD_RAF_ (0.77855 ± 0.09053); Apical proximal: Control (0.37316 ± 0.0448), H-Ras (0.84397 ± 0.0987), ΔH-Ras (0.30431 ± 0.04335), and RBD_RAF_ (0.35444 ± 0.06879); basal: Control (0.5743 ± 0.05578), H-Ras (0.95856 ± 0.06823), ΔH-Ras (0.58031 ± 0.09471), and RBD_RAF_ (0.5017 ± 0.107). Control: N=17; H-Ras: N=15; ΔH-Ras: N=10; RBD_RAF_: N=7. *p<0.05, **p<0.01, ***p<0.001 (Two-way ANOVA).

For morphometric analysis, two aspects are of importance for optimal image and analysis quality: (1) additional expression of a cell marker protein and (2) a high signal-to-noise ratio. In this study, we used a red fluorescence protein as cell marker and injected it either alone (from here on referred to as “Control”) or together with the biosensors. Sparse viral labeling via a Cre-lox approach will lead to a handful of virally transfected pyramidal cells with strong fluorescence and minimal background noise. In such manner, we generated a Cre-dependent version of the H-Ras biosensor and injected the construct that can express Cre protein in low concentration into layer 2/3 primary motor cortex (M1) (Figure 1B). Expression of the Cre protein was regulated under the CaMKII promoter to allow for pyramidal cell specificity ^42^. H-Ras expression was confirmed by H-Ras antibody staining showing strong signals in cells expressing the H-Ras biosensor, but only minor endogenous expression in control conditions (Figure S1A-C). To count and morphologically characterize dendritic spines, we imaged neurons under two-photon microscope from acute brain slices. We identified each individual spine from the selected areas on the dendritic tree letting a semiautomated, custom-made program calculate spine count as well as spine neck length. Average spine numbers were significantly higher in pyramidal neurons that expressed H-Ras (> 40%) (Figures 1C-D). Spine numbers between controls were not significantly different and were in line with previously reported spine densities ^8, 43, 44^.

Furthermore, the phenotype shifted towards spines with overall longer spine necks (Figure S2A-C). When classified into filopodia and spines, we uncovered that the overall percentage of filopodia increased, but they did not account for the total increase of all protrusions (Figure S2A-C). To determine if the effect is universal or limited to certain regions of the dendritic tree, we separated the neuron into representative dendritic segments (apical proximal, apical distal and basal) and assessed changes in spine distribution based on dendrite classes. The most pronounced effect of H-Ras overexpression was observed in apical proximal dendrites (> 130%) with a smaller effect exhibited in basal dendrites (> 60%) while apical distal dendrites showed no significant change in spine number (Figures 1C and 1E). Overall, our data shows that H-Ras affects spine number and morphology in layer 2/3 PyNs of the M1.

### Spine density changes by H-Ras ectopic expression is independent of cell types

To assess whether H-Ras produces more dendritic spines by accelerating existing molecular apparatus or rather directly drives spinogenesis, we expressed H-Ras into PV-INs and VIP-INs where key excitatory synaptic proteins are expected to be absent or low ^39, 45, 46^. The Cre-dependent biosensors and cell marker were injected into M1 layer 2/3 of PV-Cre and VIP-Cre mouse lines, respectively (Figures 2A-B). Antibody staining against PV confirmed the selective targeting of PV-INs (Figures S3A-C). Spine analysis revealed an enormous increase in spine number after H-Ras overexpression exhibiting a 300% increase in PV-INs (Figures 2C-D) and a 250% increase in VIP-INs (Figure 2E-G). Basal level spine density was similar to previous reports ^39, 44^. When looking at spine number increase in relation to distance from the cell body, VIP-INs displayed an equal rise in spine number throughout the dendritic trees (Figures 2E-G). PV-INs showed a more pronounced but equally significant effect in distal (> 350%) compared to proximal dendritic segments (> 225%) (Figures 2C-D). These changes are somewhat different from changes in distal apical dendrites of PyNs where there was no significant further increase in spine numbers, suggesting that the availability of free dendritic space that can implement more dendritic spines is an important factor. We also analyzed whether these newly formed spines in INs have longer spine necks similar to changes in PyNs. Dendritic spines in H-Ras overexpressing neurons exhibited longer spine necks in PV-INs, but not in VIP-INs (Figures S4A-B). These spines with longer spine necks opens the possibility that they are immature and silent.

**Figure 2:**
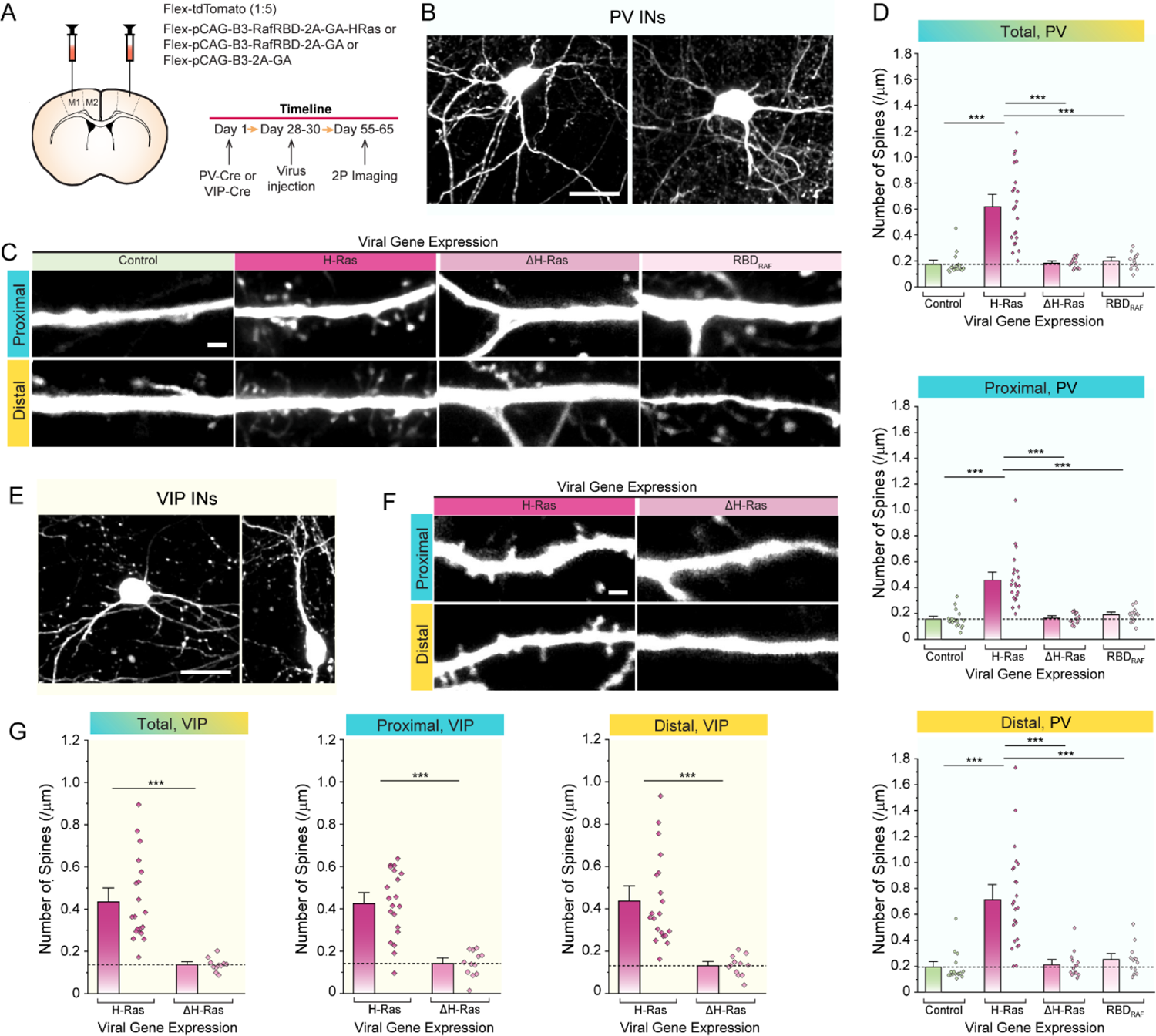
Spine density changes by H-Ras ectopic expression is independent of cell types. **(A)** Virus injection scheme and experimental timeline. **(B)** Representative two-photon microscopy images of parvalbumin-positive interneurons (PV-INs). Scale bar: 10 μm. **(C)** Two-photon microscopy images showing representative dendrites (proximal and distal) of PV-INs expressing Flex-tdTomato and either Flex-pCAG-B3-RafRBD-2A-GA-HRas (H-Ras), Flex-pCAG-B3-RafRBD-2A-GA (RBD_RAF_) or Flex-pCAG-B3-2A-GA (ΔH-Ras). Scale bar, 2 μm. **(D)** A superimposed bar and dot graph showing the spine density of each neuron (dot) and mean ± SEM (bar), respectively, for each condition (Control, H-Ras, RBD_RAF_, and ΔH-Ras). (Top) Spine density averaged from all imaged dendritic branches; Control (0.17519 ± 0.02112); H-Ras (0.61928 ± 0.06211); ΔH-Ras (0.18324 ± 0.01037); RBD_RAF_ (0.20159 ± 0.01719); (Middle) spine density of proximal dendrites; Control (0.15556 ± 0.01494); H-Ras (0.45558 ± 0.04228); ΔH-Ras (0.16542 ± 0.01075); RBD_RAF_ (0.18838 ± 0.01431); (Bottom) spine density of distal dendrites; Control (0.19191 ± 0.02918); H-Ras (0.71365 ± 0.07629); ΔH-Ras (0.21208 ± 0.02675); RBD_RAF_ (0.25007 ±. 0.03138). The dotted line represents the average spine density of dendrites expressing tdTomato only (Control). Neurons expressing H-Ras showed a higher rate of dendritic spines throughout all dendritic regions. Control: N=16; H-Ras: N=22; ΔH-Ras: N=14; RBD_RAF_: N=13. ***p<0.001 (Kruskal Wal Anova). **(E)** Representative two-photon images of vasoactive intestinal peptide-expressing interneurons (VIP-INs). Scale bar: 10 μm. **(F)** Two-photon microscopy images showing representative dendrites (apical distal, apical proximal, and basal) of VIP INs expressing Flex-tdTomato in the primary motor cortex of a typical acute brain slice made from VIP-Cre mice at ∼P60 injected 4-weeks prior with Flex-tdTomato and either H-Ras, or ΔH-Ras. Scale bar, 2 μm. **(G)** A superimposed bar and dot graph showing the spine density of each neuron (dots) and mean ± SEM (bar), respectively, for each condition (H-Ras, and ΔH-Ras). (left) Spine density averaged from all imaged dendritic branches; H-Ras (0.43456 ± 0.04397); ΔH-Ras (0.13683 ± 0.13375); (middle) spine density of proximal dendrites; H-Ras (0.42397 ± 0.0354); ΔH-Ras (0.14144 ± 0.0175); (right) spine density of distal dendrites; H-Ras (0.43697 ± 0.04737); ΔH-Ras (0.13057 ± 0.01376). The dotted line represents the average spine density of dendrites expressing ΔH-Ras. Neurons expressing H-Ras showed a higher spine density throughout all dendritic regions. H-Ras: N=20; ΔH-Ras: N=12; ***p<0.001 (Two Sample T-Test).

To sum, our data show that H-Ras increases spine numbers not only in PyNs and INs, suggesting that changes in spine number are not restricted in specific cell types. The manipulation of H-Ras expression indicates that spine formation in INs may be permissive without having the involvement of excitatory synaptic protein machinery. It also implies that the low spine density in INs may be maintained by active cellular processes preventing exuberant spine formation in normal conditions but when this system is broken, it allows a massive formation of dendritic spines.

### Local H-Ras activity increases spinogenesis

To examine if increased spine number more directly relates to H-Ras activity, we measured H-Ras activity and its correlation with the likelihood of having dendritic spines. H-Ras biosensor signals appeared dim in acute slice preparations. Additionally, the high number of spines in adult neurons, especially under H-Ras overexpression limits the analysis of spineless regions. For these reasons, we decided to express H-Ras in organotypic slice cultures during early development (EP 9-12) [EP (equivalent postnatal) day = postnatal day at slice culturing + days in vitro] where spine number is relatively low (Figures 3A-B). Similar to acute brain slices, H-Ras overexpression increased spine numbers in PyNs (Figure 3C). A DNA bullet containing H-Ras sensor and tdTomato plasmids was transfected to visualize both H-Ras activity and cell morphology from the same neuron (Figure 3D). H-Ras activity was visible along the dendrite (Figure 4E). When analyzing H-Ras activity, dendritic regions were sectioned by a 1 μm scale throughout the dendrite to have a similar size scale of dendritic spine diameter. We uncovered that H-Ras activity indeed correlates with the probability of spine appearance in a sigmoidal fashion progressing from no correlations at a low range of H-Ras activity, to a strong correlation with high H-Ras activity (Figure 3F). When registered H-Ras signals from one dendritic segment were aligned with dendritic spines in other dendritic segments, no correlation was detected (Figure 3G), suggesting that the presence of dendritic spines in the position of high H-Ras activity was not simply due to the randomly distributed high spine density. Our data show that unevenly distributed dendritic spines can be mirrored by H-Ras activity, implying the role of H-Ras in spinogenesis.

**Figure 3:**
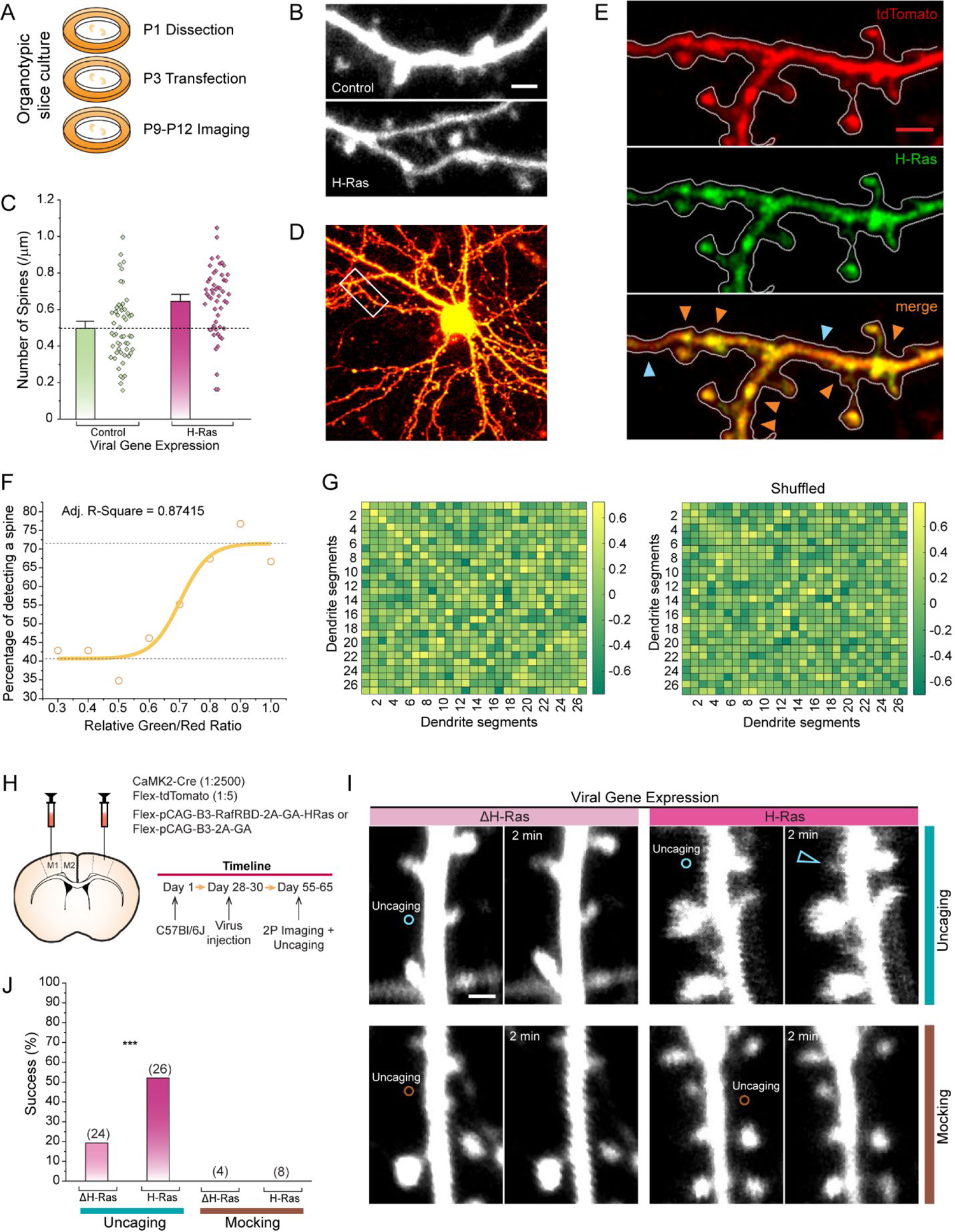
Local H-Ras activity facilitates *de novo* growth of spines. **(A)** Schematic of organotypic slice preparation and timeline. **(B)** Representative dendrite images expressing tdTomato (top) and H-Ras (tdTomato + pCAG-B3-RafRBD-2A-GA-HRas) (bottom) collected via a two-photon microscope at P12. DNA was biolistically transfected to organotypic cortical slices at P1. Scale bar: 2 μm. **(C)** A summary graph showing the spine density of each analyzed dendritic section (∼30 μm) (dots) and the mean spine density ± SEM (bar) of Control (0.49719 ± 0.02549) and H-Ras neurons (0.6445 ± 0.02608). The dotted line represents the average spine density of Control dendrites. Control: N= 53, H-Ras: N=52; ***p<0.001 (Two Sample T-Test). **(D)** tdTomato and H-Ras biosensor expression in a representative organotypic pyramidal neuron: red= tdTomato, green= H-Ras biosensor. Scale bar: **(E)** Magnification of dendritic branch shown in D (white box) (top) tdTomato, (middle) H-Ras biosensor GFP, and (bottom) merge. Colored arrow heads indicate examples of high H-Ras activity (orange) or low H-Ras activity (light blue). Scale bar: 2 μm. **(F)** Scatter plot showing that higher H-Ras activity (relative green/red ratio) correlates with a higher percentage of detecting a spine in close proximity (1 μm). The sigmoidal curve fitting to the data shows an R-Value of 0.87415. **(G)** Correlation matrix comparing H-Ras activity values with the existence of a spine in a 1 μm radius; (left) experimental data, (right) shuffled data control. H-Ras activity correlates to a higher extent with the existence of dendritic spines in the experimental data in comparison to shuffled data. (H) Virus injection scheme and experimental timeline. **(I)** Example images of high frequency glutamate uncaging (HFU) experiments (white circles, 40 pulses at 10Hz) on adult pyramidal neurons (∼P60): (top left) ΔH-Ras + MNI glutamate, (top right) H-Ras + MNI glutamate, (bottom left) ΔH-Ras without MNI-glutamate, and (bottom right) H-Ras without MNI-glutamate. All neurons expressed tdTomato as cell markers. White arrowheads indicate *de novo* spine formation after HFU. Scale bar, 2 μm. **(J)** Success rate of *de novo* spine formation by HFU at P60 in pyramidal neurons expressing tdTomato and either ΔH-Ras (19.2%) or H-Ras (52%). In the absence of MNI-glutamate (mock) neither ΔH-Ras nor H-Ras exhibited any *de novo* spine formation. ΔH-Ras: N= 24 trials, 16 cells; H-Ras: N= 26 trials, 15 cells; ΔH-Ras mock: N= 4 trials, 4 cells; H-Ras mock: N= 8 trials, 7 cells. ***p<0.001 (Chi-Square Test).

**Figure 4:**
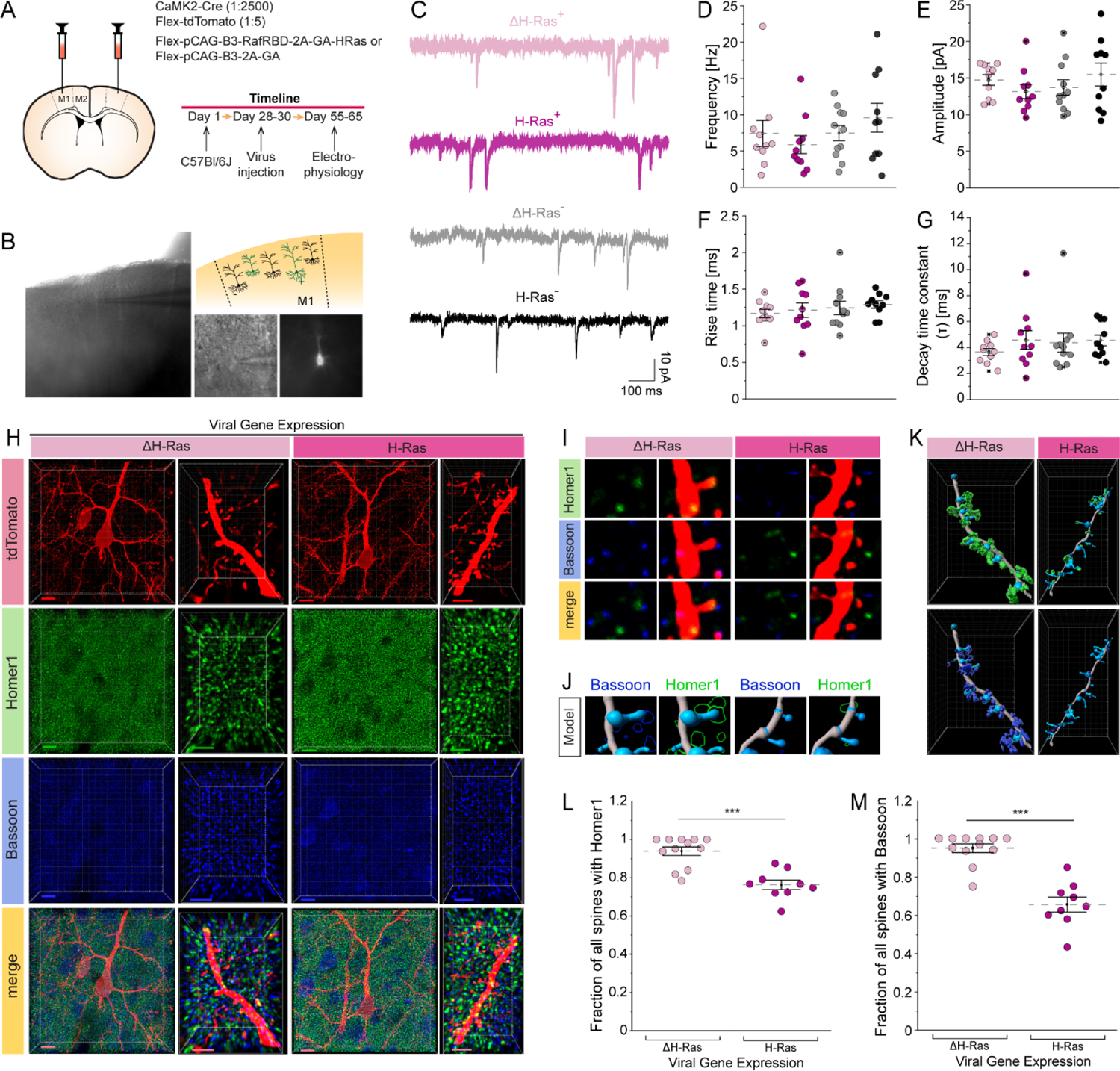
Increased spine number does not represent features of functional excitatory synapses. **(A)** Virus injection scheme and experimental timeline. **(B)** Light microscopy images of layer 2/3 pyramidal neuron recorded in whole-cell voltage-clamp-mode in acute slices: whole motor cortical brain region (left), magnified image: light microscopy and fluorescent (right bottom), schematic of pyramidal neurons being fluorescent positive (+) or fluorescent negative (-) (right top). **(C)** Examples traces of mEPSCs in ΔH-Ras^+^ (blue), H-Ras^+^ (orange), ΔH-Ras^-^ (grey), and H-Ras^-^ neurons (black). **(D-G)** mEPSC frequencies (D), amplitudes (E), rise times (F), and decay times (G) in ΔH-Ras^+^ (blue), H-Ras^+^ (orange), ΔH-Ras^-^ (grey), and H-Ras^-^ neurons (black). Each dot represents an average value from one neuron, and the dotted line represents mean ± SEM. **(H)** Example confocal images of ΔH-Ras (left) and H-Ras of pyramidal neurons and magnified dendrites (right). From top to bottom: image of pyramidal neuron expressing tdTomato, postsynaptic protein Homer1 stained with Alexa Fluor 633 (green), presynaptic protein bassoon stained with Alexa Fluor 405 (blue), merge of all channels. Scale bars: 10 μm (pyramidal neuron), 3 μm (dendrite). **(I)** Magnification image of (H) visualizing single spines of ΔH-Ras (left) and H-Ras (right). From top to bottom: postsynaptic protein Homer1 (green) stained with Alexa Fluor 633 with and without dendrite visualization, presynaptic protein bassoon with Alexa Fluor 405 (blue) stained with Alexa Fluor 405 with and without dendrite visualization, merge of channels with and without dendrite visualization. Scale bar: 0.75 μm. **(J)** Imaris reconstructed image of dendrites and dendritic spines in conjunction with Bassoon reconstruction (blue) or Homer1 reconstruction (green) of ΔH-Ras (left) and H-Ras neurons (right). Scale bar: 0.75 μm. **(K)** Imaris reconstruction of dendrites/spines together with Homer1 (green, top) or bassoon (blue, bottom) of ΔH-Ras (left) and H-Ras neurons (right). **(L-M)** Analysis of fraction of spines encompassing Homer1 puncta (L) or having close contact (0.1 μm) to bassoon puncta (M) in either ΔH-Ras or H-Ras neurons shows that neurons expressing ectopic H-Ras have reduced number of spines expressing Homer1 as well as reduced bassoon contacts. ΔH-Ras: N= 12, H-Ras: N= 9; **p<0.01, ***p<0.001 (Two-Sample Kolmogorov-Smirnov Test).

To find additional evidence showing the involvement of H-Ras in spine growth, we performed two-photon glutamate uncaging experiments ^14–16, 18, 47, 48^. A mixture of AAV expressing CaMKII-Cre (1:2,500 dilution), Flex-tdTomato, and Flex-pCAG-B3-RafRBD-2A-GA-H-Ras or Flex-pCAG-B3-2A-GA were injected in layer 2/3 neurons of the M1 and acute brain slices were made one month after viral injection (Figure 3H). We perfused 3 mM (4-methoxy-7-nitroindolinyl)-glutamate (MNI-Glu) into the recording chamber containing Mg^2+^-free ACSF. A brief pulse of 720 nm laser (30 pulses at 10 Hz, 4 ms duration) was delivered 0.5 μm away from the dendrite (Figure 3I). We found that glutamate uncaging significantly increased the success rate of *de novo* spine growth when H-Ras was transfected (Figure 3J). The spine growth was dependent on glutamate release because the same laser pulses without MNI-Glu failed to induce spine growth (Figure 3J). Thus, our data strongly suggest that high H-Ras mediated the formation of new dendritic spine growth.

### H-Ras increases spine dynamics

Increased spine density could be achieved not only by more spine formation but also by increased stability. To examine the spine stability, we examined spine turnover rates in layer 2/3 PyNs. We performed time lapse-imaging of dendrites (Figure S5A). In a condition where H-Ras was ectopically expressed, the percentage of stable spines decreased with higher morphological changes. The number of newly formed spines were higher than eliminated ones, suggesting that H-Ras plays a major role in producing more dendritic spines rather than making existing spines more stable (Figure S5B).

### Increased spine number does not represent features of functional excitatory synapses

We next sought to determine whether the exuberant number of spines following H-Ras overexpression are coupled with functional synaptic changes. To test functional connectivity changes, we measured miniature excitatory postsynaptic currents (mEPSCs) in layer 2/3 PyNs of M1 (Figures 4A-B). Changes in mEPSC frequency are thought to reflect a shift in functional spine number ^49^. In our case, an increase would correlate with a rise in functional spines, while a reduced or unchanged mEPSC frequency would point towards silent, non-functional spines. *Post-hoc* blind analysis showed that there was no difference in mEPSC frequency, amplitude, rise or decay time between control and H-Ras conditions (Figures 4C-G), arguing against the notion that dendritic spines should always be considered functional excitatory synapses. This result seems to be in line with recent studies demonstrating the existence of silent filopodia in the adult neocortex ^25, 50^.

We also examined whether dendritic spines possessed key pre- and post-synaptic proteins normally found in excitatory synapses. We used antibodies against the postsynaptic scaffolding protein Homer1, which is involved in glutamate receptor targeting. Homer1 comes in three isoforms with its two most abundant isoforms exclusively expressed in the postsynaptic density (PSD), with the presence of a PSD being a hallmark for mature spines ^51–53^. Image analysis was performed by identifying spines colocalized with Homer1 in first-order dendritic branches (Figure 4H). Since we could not distinguish between normal, preexisting spines and new H-Ras-induced spines, we selected the entire spine population to calculate the fraction of spines containing Homer1. The number of spines with a Homer1 surface contact and the number of spines with a Bassoon surface contact were recorded. The presence of Homer1 or Bassoon signal close to the spine head was counted as colocalization (within 0 μm from the spine head for Homer1 and 0.1 μm for Bassoon) (Figure 4I-J). We found that significantly less spines contained Homer1 in H-Ras transfected neurons (Figure 4K-M), suggesting spines lacking Homer1 staining were induced by H-Ras ectopic expression. Previously reported silent spines lack AMPA (α-amino-3-hydroxy-5-methyl-4-isoxazoleproprionic acid) type receptors but show normal presynaptic connectivity ^25, 54, 55^. We checked if H-Ras induced spines form presynaptic networks even though they are postsynaptically immature. Consistent with Homer1 data, the fraction of spines innervated by Bassoon was significantly lower with H-Ras overexpression suggesting that H-Ras induced spines fail to build connections with presynaptic partners.

## DISCUSSION

It has been long belief that dendritic spines are structural correlates of excitatory synapses ^56–58^. Because this concept has been confirmed by numerous functional and anatomical studies, evidence against this notion has been hardly found. When H-Ras was overexpressed in layer 2/3 PyNs, the number of dendritic spines was largely increased. The increase of spine number was observed even in proximal apical dendrites where the number of excitatory synapses is relatively lower. The most striking changes are found in INs. When H-Ras was ectopically expressed in INs, the increase was not just 20% or 30%, but twice or three times higher than the spine number in control conditions. Such uncommon changes implied that H-Ras induced dendritic spines may not represent the preexisting spine group playing a role as functional excitatory synapses. The morphology of dendritic spines in H-Ras expressing neurons also looked like a filopodium with longer spine neck.

Molecules involved in dendritic spine formation in INs have been poorly identified. One study reported that the overexpression of extracellular N-terminal domain of AMPA receptor subunit GluA2 increased spine number in GAD-positive neurons ^59^. However, experiments were only performed in embryonic culture neurons with demonstration of a few sample images. This finding is also not compatible with a more recent finding where spine density was completely normal even when all subunits of AMPARs and N-methyl-D-aspartate receptors (NMDARs) were deleted ^23^. Therefore, mechanistic understanding of spine formation in INs has still not been understood, and this study presents the permissive processes of spine outgrowth in both PyNs and INs. The capability of having a high number of dendritic spines in INs implies that the reason for maintaining the low density of dendritic spines in INs could be due to active precluding mechanisms. One promising candidate as preclusion molecule is the paired Ig-like receptor B (PirB), with its autonomous nature to limit spine density when expressed in L2/3 PyNs of the visual cortex ^60^. However, when exploring the differential mRNA expression of PirB across various neuron subtypes using the searchable web interface that accesses transcriptomic datasets (http://research-pub.gene.com/NeuronSubtypeTranscriptomes) ^46^, we noticed that PirB expression levels were highest in PyNs, with only minor expression in PV and VIP INs. This suggests that PirB may not play a factor in the preclusion of dendritic spines in INs.

The dendritic region with high H-Ras activity displayed higher probability to detect dendritic spines. The H-Ras biosensor allows changes in fluorescence, so we were able to detect correlation between H-Ras activity and spine formation. Because H-Ras is freely diffusible along the membrane, it is likely that the correlation between spine density and H-Ras activity was due to the presence of high H-Ras activity in a dendritic region with high surface area to volume ratio. More direct evidence of H-Ras involvement in spinogenesis was confirmed by two-photon glutamate uncaging experiments. Higher success rate of *de novo* spine formation by glutamate photolysis suggest that H-Ras overexpression increased the sensitivity of downstream signaling of NMDARs. Spine turnover rates confirmed the involvement of H-Ras in spine formation, rather than stabilization, as there was a significant increase in spine formation with little change in spine elimination. Although spines exist in a wide range of sizes and shapes following a continuum rather than segregated classes, immature spines are thought to exhibit elongated necks and thin heads resembling filopodia ^61–63^. In agreement, our data showed an overall shift to significantly longer spine necks in PyNs and PV-INs, but not VIP-INs, after H-Ras overexpression. Upon categorizing protrusions as either filopodia or spines, we observed that the proportion of filopodia increased.

Our data further show that the exuberant spines were not functionally incorporated into the pre-existing neural network. Upregulation of functional spines alters physiological properties of neurons, with more spines leading to a higher frequency of miniature excitatory postsynaptic currents (mEPSCs) ^49^. This change in mEPSC frequency was absent in H-Ras expressing layer 2/3 PyNs of M1, indicating that the exuberant spines are silent or not even making contact with presynaptic neurons. Another explanation is that filopodia produced by H-Ras are silent, and still waiting for becoming mature, functional dendritic spines at the time of data analysis, but this scenario is unlikely because we recorded neurons about one month after H-Ras was expressed. Supporting this interpretation, spines in H-Ras transfected neurons were devoid of a key postsynaptic molecule, reflected by the absence of Homer1 in ∼25% of spines. Even though we are unable to distinguish between H-Ras induced and pre-existing spines, the fraction of spines without Homer1 was comparable to that of new spines. Homer1a is the only isotype among the three types of Homer1 not bound to the PSD, and is instead present in the cytosol and found in both mature and immature spines ^51^, which may explain the small discrepancy observed between the total spine increase and the number of spines lacking Homer1. These spines are still incorporated into the synaptic network and have a presynaptic partner although they are functionally silent. In contrast, our antibody staining against the presynaptic marker bassoon revealed that about 35% of spines (comparable with the 40% spine increase) were without bassoon signals after H-Ras overexpression.

Since the dendritic spines formed via ectopic H-Ras expression seem to not build functional synapses, at least not immediately, this opens the door for speculations. Could these protrusions have another purpose? In certain cell types such as PC12 cells, HeLa cells, microglia, astrocytes and pericytes, intercellular connections can form through structures known as tunneling nanotubes, through which the cells can exchange nutrients, and small molecules ^64–69^. These connections were shown to arise from filopodia outgrowths where two filopodia from two different cells form a filopodia bridge ^69^. Tunneling nanotubes formed between B and T immune cells where enriched for H-Ras suggesting that H-Ras may be transported through these nanotubes as cargo or is involved in their buildup ^70^. Although tunneling nanotubes have not been described in neurons, the abundance of filopodia in these cells suggests that similar intercellular communication structures may exist. Hence, we hypothesize that H-Ras may play a key role in the formation of these tunneling nanotubes, potentially contributing to intercellular communication in neuronal networks. More research is needed to investigate this hypothesis and unravel the precise mechanisms involved.

Our data uncovered that dendritic spines can be induced without causing functional synaptic connectivity changes. The spine-synapse dissociation and robust increase of dendritic spines in INs suggest that dendritic spines may not be just created to be connected with other neurons. There might be a crossroad where the fate of these protrusions is decided, developing into dendritic spines or other structures like tunneling nanotubes.

### LIMITATIONS OF THE STUDY

We used two-photon microscopy in our study. However, recent studies combining light and electron microscopy have shed light on some of the limitations of conventional light microscopy in capturing small, thin structures such as filopodia, which may be obscured by the fluorescence surrounding dendrites and larger spines ^25, 27, 50^. As a result, the generation and subsequent transition of certain filopodia into spines may not be observable using our current methods.

More rigorous microscopy techniques, such as electron microscopy, may be necessary to observe these structures accurately. Unfortunately, timelapse imaging and high-frequency uncaging cannot be deciphered with electron microscopy. Furthermore, the precise molecular mechanisms of how H-Ras induces *de novo* spines were not the focus of this study and need to be investigated in follow-up studies.

## AUTHOR CONTRIBUTIONS

S.K. and H-B.K. conceived and designed the study. S.K. and I.D. performed viral injection for experiments. S.K prepared organotypic slices. S.K. performed two-photon imaging, two-photon uncaging, and electrophysiology recording. S.K. and I.D performed immunohistochemistry and confocal imaging. M.C. wrote spine morphology data analysis and deconvolution program. S.K wrote correlation analysis program. S.K. analyzed spine density, morphology, dynamics, correlation, and electrophysiology data. I.D. provided assistance in spine morphology analysis. I.D and S.K analyzed histology data. S.S provided Cre-dependent versions of the H-Ras biosensor. H-K.L. and W-D.H provided critical advice on the manuscript. S.K. and H-B.K. wrote the manuscript, and S.K. and H-B.K. edited the manuscript. All authors discussed and commented on the manuscript.

## ACKNOWLEDGEMENTS

We thank members of the Kwon laboratory for helpful discussions. We thank Patricia Janak, Marshall Schuler, and Seth Margolis for comments on the experiments. We thank Mary McMillan for helping in experiment sample preparation, and we thank the Kanold laboratory for help with the initial electrophysiological experiments. This work was supported by Johns Hopkins School of Medicine (to H-B.K.), Max Planck Florida Institute for Neuroscience (to H-B.K.), the National Institutes of Health Grants DP1MH119428 (to H-B.K), the KAIST-funded Global Singularity Research Program for 2022 (W.D.H.).

## STAR METHODS

### CONTACT FOR REAGENT AND RESOURCE SHARING

Further information and request for resources or analysis code should be directed to and will be fulfilled by the Lead Contact, Hyung-Bae Kwon.

### EXPERIMENTAL MODEL AND SUBJECT DETAILS

#### Animals

C57BL/6 (Cat. #: 000664, 4-9 weeks old), PV-Cre (Cat.#:017320, 4-9 weeks old) and VIP-Cre mice (Cat.#:010908, 4-9 weeks old) from Jackson laboratory (Bar Harbor, ME, USA) were used for experiments. All mice were maintained in a temperature and humidity-controlled facility following a 12 hr light/ 12 hr dark cycle with food and water available ad libium. Control and test group animals were randomly chosen, and similar numbers of male and female mice were used for experiments. All experimental procedures were carried out in accordance with protocols approved by Johns Hopkins University Animal Care and Use Committee, the Max Planck Florida Institute for Neuroscience Institutional Animal Care and Use Committee, and National Institutes of Health guidelines.

#### Tissue culture

Newborn C57BL/6 pups (1-2 days postnatally) were prepared for organotypic hippocampal slice cultures according to published procedure ^71^. Preparations were executed in an aseptic environment. Pups were decapitated, the brains were surgically removed and placed into ice-cold dissection medium (in mM: 1 CaCl_2_, 5 MgCl_2_, 10 D-Glucose, 4 KCl, 26 NaHCO_3_, 234 Sucrose). Each hemisphere’s cortices were isolated and transferred onto a tissue chopper stage in a sterile preparation hood. Any excess liquid around the tissue was removed before slicing the cortices into 300 μm thick coronal sections. For easier handling, slices were transferred to a plastic dish containing culture medium (in mM: 1 L-Glutamine, 1 CaCl_2_, 2 MgSO_4_, 13 D-Glucose, 5.3 NaHCO_3_, 30 Hepes; 8.4 g/l MEM Eagle medium, 1mg/l Insulin; 20% Horse serum, 0.00125% Ascorbic Acid) and then placed one-by-one onto small membrane cell culture inserts (Millipore, Burlington, MD). Maximally three cortical slices were cultured in each cell culture insert. Slices were maintained in culture medium for up to 2 weeks at 37°C/5% CO_2_ changing 70% of the medium every 2-3 days. Slices were transfected 1 day after preparation using biolistic gene transfer as described previously ^72^. A total of 10 μg of tdTomato alone or together with 30 μg pCAG-B3-RafRBD-2A-GA-HRas were coated onto 6-7 mg of gold particles. Slices were imaged 7-11 days after transfection.

## METHODS DETAILS

### Animal surgery and viral stereotaxic injection

Surgeries were conducted on ∼4-week-old mice in aseptic conditions using a small animal stereotactic setup (Kopf instruments, Tujunga, CA, USA). Mice were fully anesthetized in a closed chamber with isoflurane (5%) before moving animals to the stereotaxic setup where the isoflurane concentration was reduced to 2% and the mice’s head was fixed. Ophthalmic ointment (Puralube Vet Ophthalmic Ointment) was applied to both eyes to prevent drying and body temperature (37 °C) was maintained by a thermostatically controlled heating pad (Harvard Apparatus, Holliston, MA, USA). To reduce the infection risk, mice’s hair was removed with hair removal lotion (Nair, Church & Dwight) and the exposed scalp was subsequently disinfected three times alternately with 10% betadine solution (Purdue product LP, Stamford, CT, USA) and 70% ethanol wipes. A minor incision of the scalp was made, and the periosteum removed to allow for a small craniotomy (∼0.5 mm in diameter) over the injection site. To ensure the correct placement of the craniotomy lambda and bregma lines were used for orientation.

The coordinates of the motor cortex for viral injection were anteroposterior (AP) +0.25 mm from bregma, mediolateral (ML) +/-1.5 mm from bregma, and dorsoventral (DV) −0.3 mm from the brain surface. The viral constructs (0.5 ul per hemisphere) were injected via a beveled glass micropipette (tip size 10–20 μm diameter, Braubrand), backfilled with mineral oil. Flow rate (150-250 nl/min) was regulated by a syringe pump (World Precision Instruments). The glass micropipette was fixed in place for 5 additional minutes after injection ended to minimize backflow. Finally, the scalp was stitched using vetbond glue (3M Animal care products, St. Paul, MN, USA). General analgesia (Buprenorphine SR, 0.6 mg/kg, or Meloxicam, 0125 mg) was injected subcutaneously or given orally, and mice were monitored until they recovered from anesthesia.

### Preparation of acute cortical slices

Surgery animals recovered for 4 weeks after viral injection before they were anesthetized with isoflurane and either directly decapitated (for two-photon imaging) or cardiac perfused with ice-old cutting solution (in mM: 215 Sucrose, 20 D-Glucose, 26 NaHCO_3_, 4 MgSO_4_, 4 MgCl, 1.6 NaH_2_PO_4_, 1 CaCl_2_, 2.5 KCl) prior to decapitation (for electrophysiology). The brains were quickly removed and submerged in ice-cold cutting solution. Cortical slices (300 μm thick) were prepared using a VT1000S vibrating microtome (Leica) and then incubated at 30°C for 30 minutes in a holding chamber filled with artificial cerebrospinal fluid (ACSF) (in mM: 124 NaCl, 3 KCl, 1.3 MgSO_4,_ 2.5 CaCl_2_, 10 D-Glucose, 1.25 NaH_2_PO_4_, NaHCO_3_). The holding chamber was moved to room temperature and the slices recovered for 30 more minutes before imaging or recording. All solutions were saturated for at least 30 min with 5% CO_2_/95% O_2_.

### Two-photon imaging

Two-photon imaging was performed on transfected layer 2/3 pyramidal neurons from both organotypic and acute slices at either 10-14 days in vitro (DIV) or on postnatal days 58-62, respectively. Layer 2/3 VIP and PV interneurons were only studied in the acute slice imaging context. Slices were placed into a submersion-type chamber (Warner Instruments, Holliston, MA, USA) containing recirculating, oxygenated ACSF (in mM: 124 NaCl, 3 KCl, 1.3 MgSO_4,_ 2.5 CaCl_2_, 10 D-Glucose, 1.25 NaH_2_PO_4_, NaHCO_3_). Selected neurons were within 40 μm of the slice surface and image stacks (512 × 512 pixels; 0.049 μm/pixel) with 0.5 to 1-μm z-steps were collected from proximal apical (<50 μm from soma), distal apical (>50 μm from soma) and basal dendrites for pyramidal cells and from proximal (<50 μm from soma) and distal (>50 μm from soma) dendrites for VIP and PV interneurons using a two-photon microscope (Prairie Technologies, Inc) with a pulsed Ti::sapphire laser (MaiTai HP DeepSee, Spectra Physics) tuned to 920 nm (5-7.5 mW at the sample) under a 60x objective (1.0 NA, Olympus). For spine dynamics assessments in pyramidal cells, images of ∼4 dendritic regions per neuron were taken every 10 minutes for a period of 1 hour. All images shown are maximum projections of 3D image stacks after applying a median filter (2 × 2) to the raw image data.

### Glutamate uncaging

Two-photon uncaging of MNI-glutamate was implemented, as previously described ^15^. Two pulsed Ti:Sapphire lasers (Chameleon, Coherent, Santa Clara, CA) were used for imaging and uncaging with wavelengths of 920 nm and 720 nm, respectively. For MNI-glutamate uncaging on layer 2/3 pyramidal cells, 5 mM MNI-caged-glutamate was perfused into the slice chamber with existing, recirculating Mg^2+^-free ACSF (in mM: 124 NaCl, 3 KCl, 0 MgSO_4,_ 2.5 CaCl_2_, 10 D-Glucose, 1.25 NaH_2_PO_4_, NaHCO_3_), and ∼30 mW of 720 nm light at the back aperture of the objective (60x 1.0 NA objective, Olympus) was used to release the uncaging group. Glutamate high-frequency uncaging (HFU) stimuli consisted of 30 pulses (720 nm, 10-15 mW at the sample) with a duration of 4 ms delivered at 10 Hz. Uncaging locations were manually positioned in close vicinity (<0.5 μm) from the edge of the dendrite. The selected areas were well isolated with a smooth outer membrane and had at least one neighboring spine within 5 µm to ensure competency for spinogenesis. No more than four spinogenesis trials were performed from the same neuron. The mock stimulus was identical in parameters to the HFU stimulus, except carried out in the absence of caged compounds. Image stacks (512 × 512 pixels; 0.033 μm/pixel) with 1-μm z-steps were collected immediately before and after (<1 minute) HFU as well as 2, 4, 6, 10 and 15 minutes after HFU to determine the time course of spine generation. If a new spine formed within the first 5 minutes after HFU, uncaging was denoted as success. Mock uncaging trials and observation of fluorescent changes of regions adjacent to uncaging spots were used as control.

### Electrophysiology

Acute brain slices were transferred to a submersion-type, temperature-controlled recording chamber perfused with oxygenated ACSF (in mM: 124 NaCl, 3 KCl, 1.3 MgSO_4,_ 2.5 CaCl_2_, 10 D-Glucose, 1.25 NaH_2_PO_4_, NaHCO_3_) containing TTX (1 μM), D-APV (100 μM) and bicculine (20 μM). Whole-cell recordings were performed at 30°C ± 2°C on visually identified and transfected layer 2/3 pyramidal neurons of the motor cortex using a MultiClamp 700B amplifier (Molecular Devices) and an upright microscope (E600 FN, Nikon) with oblique infrared and fluorescent illumination. Neurons were patched in voltage-clamp configuration (V_hold_ = −65 mV) using borosilicate glass pipettes (electrode resistances 3-7 MΩ) filled with internal solution (in mM: 120 CsOH, 120 Gluconic acid, 10 phosphocreatine, 0.5 GTP, 4 ATP, 8 KCl, 1 EGTA,10 HEPES and 5 QX-314). Data was acquired with a custom script in Igor Pro software (WaveMetrics).

Trace duration was 10 seconds, and a test pulse was given at the beginning of each trace. Only traces with a stable steady-state holding current (<200 pA), a series resistance between 20-40 MΩ and an input resistance (R_i_) > 150 MΩ were considered for further analysis. Additionally, if either the series or input resistance shifted by more than 15% from the average, the corresponding trace was excluded. Analysis of mEPSCs events was performed using a custom made MatLab script (provided by Bryce Grier) and a minimum of 200 isolated events from each cell were quantified.

### Immunohistochemistry and confocal imaging

On ∼P55-P65, mice were deeply anesthetized with isoflurane and cardiac perfused with phosphate-buffered saline (PBS), followed by 4% paraformaldehyde (PFA). Brains were then collected and post-fixed in PFA for 24 hours. Using a VT1000S vibratome (Leica Biosystems, Buffalo Grove, IL, USA) brains were coronally sectioned into 40 μm slices. Motor cortex sections were isolated using landmarks and neuroanatomical nomenclature in accordance with the mouse brain atlas ^73^. Slices were then washed for 30 minutes with 50% ethanol, followed by a 1-hour blocking step with 10% normal goat serum in PBS and 0.2% Triton-X100. Primary antibodies (Homer1, 1:500, #160 006, Synaptic systems; Bassoon, 1:300, # 141 004, Synaptic systems; H-Ras, 1:30, # 18295-1-AP, Proteintech; Parvalbumin, # 235, Swant, 1:500) were incubated on sections for 3 days at 4°C. Subsequently, sections were rinsed three times with PBS and incubated with species-appropriate secondary antibodies for 3 h (Goat anti-guinea pig IgG Alexa Fluor 405, Abcam, ab175678; Goat anti-rabbit IgG Alexa Fluor 633, Thermo Fisher Scientific, A-11036, Goat anti-mouse IgG Alexa Fluor 633, Thermo Fisher Scientific, A-21050). After another round of three PBS washes, slices were mounted onto microscope slides (VWR, Atlanta, GA) using mounting product (ProLong™ Glass Antifade Mountant, ThermoFisher) and stored at a dark and dry place to polymerize for at least 24 hours before imaging. Images of layer 2/3 pyramidal neurons in motor cortex were acquired using an upright laser scanning confocal microscope (Zeiss LSM800, Germany) with 2.5x or 63x oil objectives. To improve signal-to-noise ratios, images were deconvolved via a custom-made MATLAB script.

## QUANTIFICATION AND STATISTICAL ANALYSIS

### Structural imaging analysis

Spine numbers and morphology of two-photon images were determined using a custom-made MatLab Script. For analysis, spines were manually selected by visual cues. Each spine was marked in the center of the spine head (highest fluorescent intensity) and at the spine origin on the dendrite. Distances between spine origin and spine head center were calculated as an estimate of real spine neck length. Next, to assess the total spine number of a neuron or different dendritic segments of a neuron (apical proximal, apical distal or basal) the sum of the detected spines was divided by the total dendrite length of each sector. For spine dynamic analysis, timelapse images were carefully investigated marking spines which changed their appearance between timepoints manually. The ratio of newly formed or eliminated spines per dendrite per neuron was compared.

### Immunohistochemistry image analysis

Imaris 9.9.1 was used to analyze and compare the antibody staining patterns of Homer1 and Bassoon between the H-Ras and ΔH-Ras (control) neurons, which were initially blinded. For each neuron imaged, 2-3 dendritic sections with clearly visible spines were cropped and used for analysis. Dendrite diameters and spines, shown by the red channel, were reconstructed manually using the filament tool. Dendrite length and spine number were recorded. Spines were then isolated as separate filaments and a new gray channel was created to represent them, which was then used to create spine surfaces. Next, the homer1 puncta, represented by the far red channel, and the bassoon puncta, represented by the blue channel, were reconstructed as surfaces. Homer1 within 0 μm of the spine surfaces and Bassoon within 0.1 μm of the spine surfaces (measurement taken from edge of one surface to edge of the other surface) were isolated as new surfaces. The number of spines with a Homer1 surface contact and the number of spines with a Bassoon surface contact were recorded. Following the Imaris analysis, neurons were unblind.

### Statistics

All statistical analyses were executed with Origin 2020b (OriginLab Corp., Northampton, MA). Probability distributions were compared using the Kolmogorov-Smirnov test and differences in variance were tested via a Levene test. Unless otherwise stated, parametric data were carried out by t test or one-way ANOVA followed by the Sidak, Holm’s, or Tukey post hoc analysis for comparisons of interactions, respectively. Non-parametric data were analyzed by Kruskal-Wallis one-way analysis of variance on ranks followed by the Dunn post hoc analysis for comparisons of interactions. Success rate of de novo spinogenesis was compared by two-sided Fisher’s exact test. Exact statistical details of experiments are noted in the figure legends. p values < 0.05 were considered statistically significant. Data are presented as mean ± SEM.

## SUPPLEMENTAL INFORMATION

**Figure S1:**
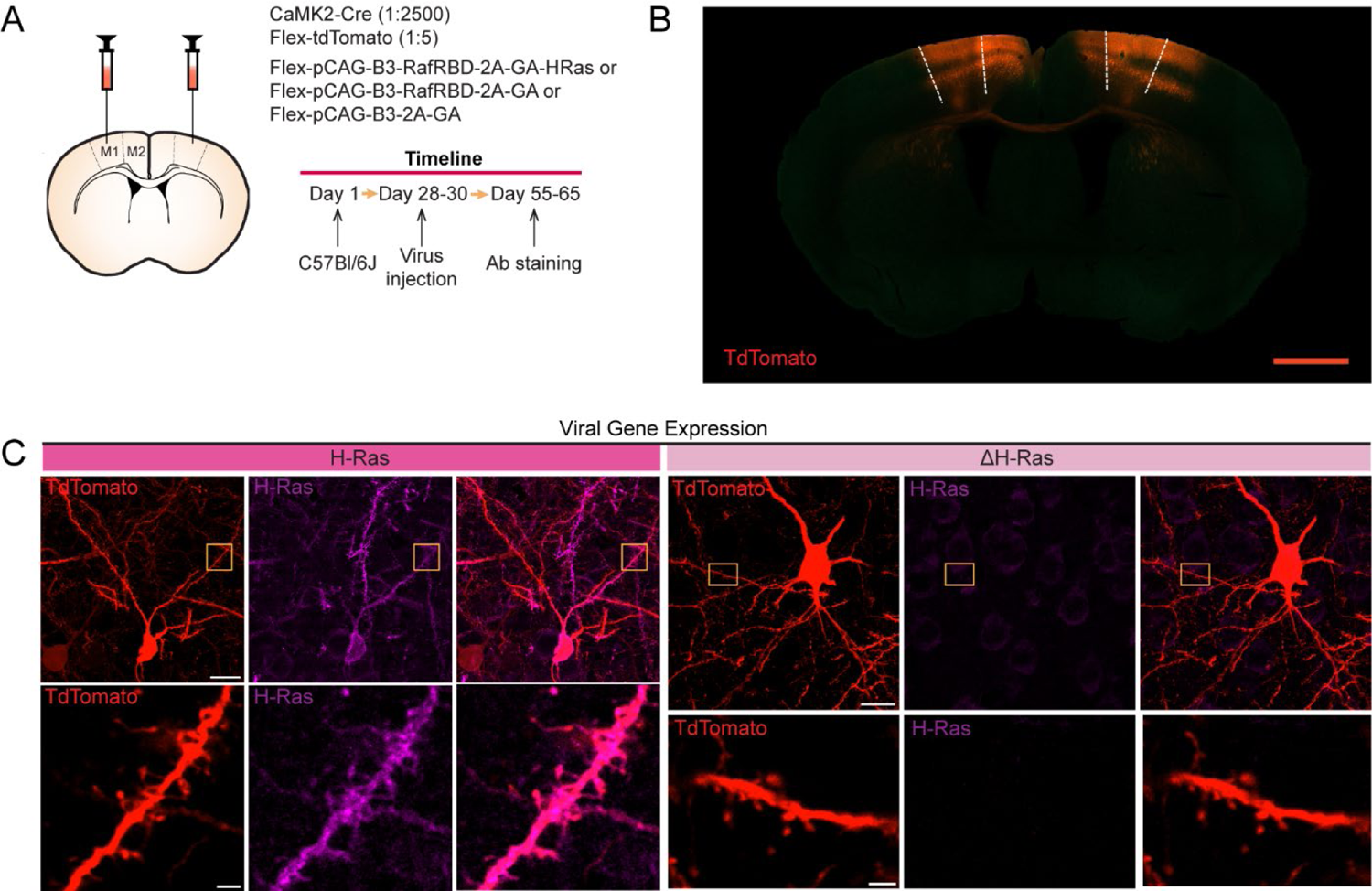
Ectopic expression of H-Ras. **(a)** Virus injection scheme and experimental timeline. **(b)** An example confocal overview image of tdTomato and ΔH-Ras expressing pyramidal cells at age ∼P60 of a 40 μm fixed coronal brain slice. Scale bar: 1 mm. **(c)** Representative confocal image of pyramidal neurons (top) and magnified dendrites (bottom) of H-Ras (left) or a ΔH-Ras expressing cells (right). Images within the yellow box was zoomed in. From left to right: Neuron and dendrite images expressing the tdTomato, H-Ras protein stained with Alexa Fluor 633 (purple), and merge of channels. Scale bar: 10 μm (top), 2 μm (bottom).

**Figure S2:**
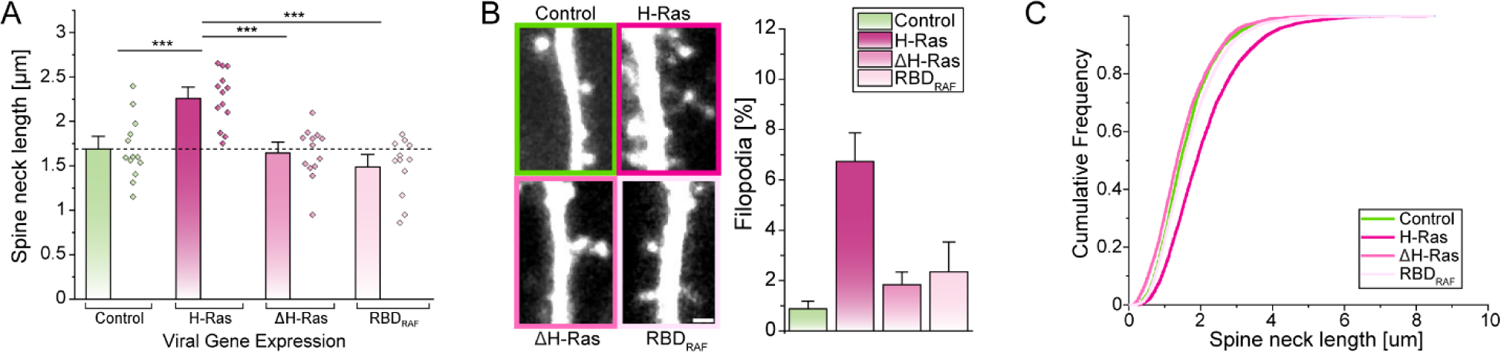
Increased spine neck length in H-Ras induced dendritic spines. **(a)** A summary graph showing the spine neck length of individual neurons (dots) and the mean ± SEM spine neck length (bar). Color scheme: Control (green), H-Ras (magenta), ΔH-Ras (light magenta), and RBD_RAF_ (pink). **(b)** Representative two-photon images of dendritic spines and their morphology. The percentage of filopodia at each expression condition. Pyramidal neurons expressing ectopic H-Ras have higher percentages of filopodia. **(c)** A cumulative frequency plot showing the entire distribution of spine neck length. Control: N= 13, H-Ras: N= 13, ΔH-Ras: N= 15, and RBD_RAF_: N= 10. ***P<0.001.

**Figure S3:**
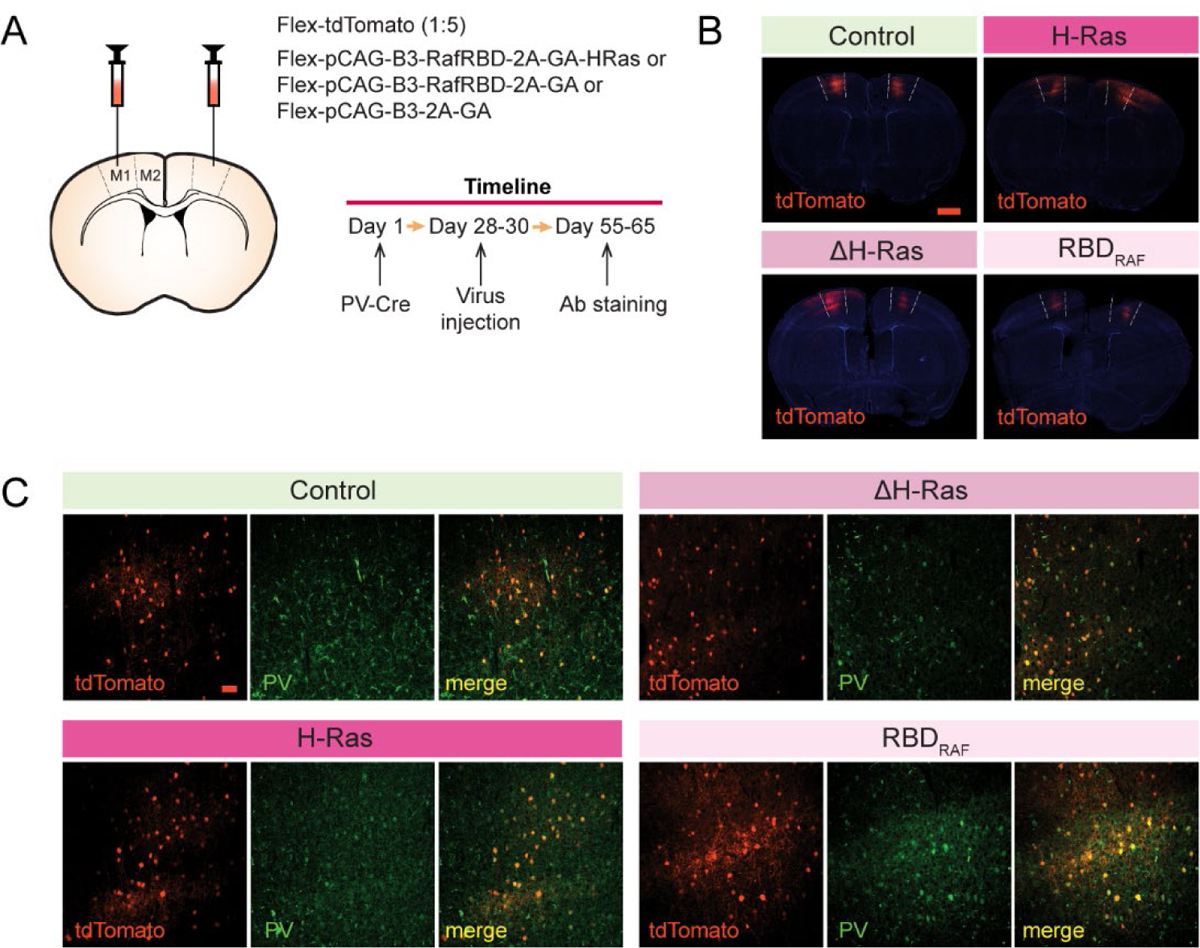
Ectopic expression of H-Ras in PV-positive interneurons. **(a)** Virus injection scheme and experimental timeline. **(b)** Example confocal images of pyramidal neurons expressing tdTomato (Control, top left) and either H-Ras (top right), ΔH-Ras (bottom left), or RBD_RAF_ (bottom right) at age ∼P60 of a 40 μm fixed coronal brain slice. Scale bar: 1 mm. **(c)** Representative confocal images showing that virus injected cells (red) are PV positive (green) stained with Alexa Fluor 633 no matter if they express tdTomato only (Control) (top left), ΔH-Ras (top right), H-Ras (bottom left), or RBD_RAF_ (bottom right). Scale bar: 50 μm.

**Figure S4:**
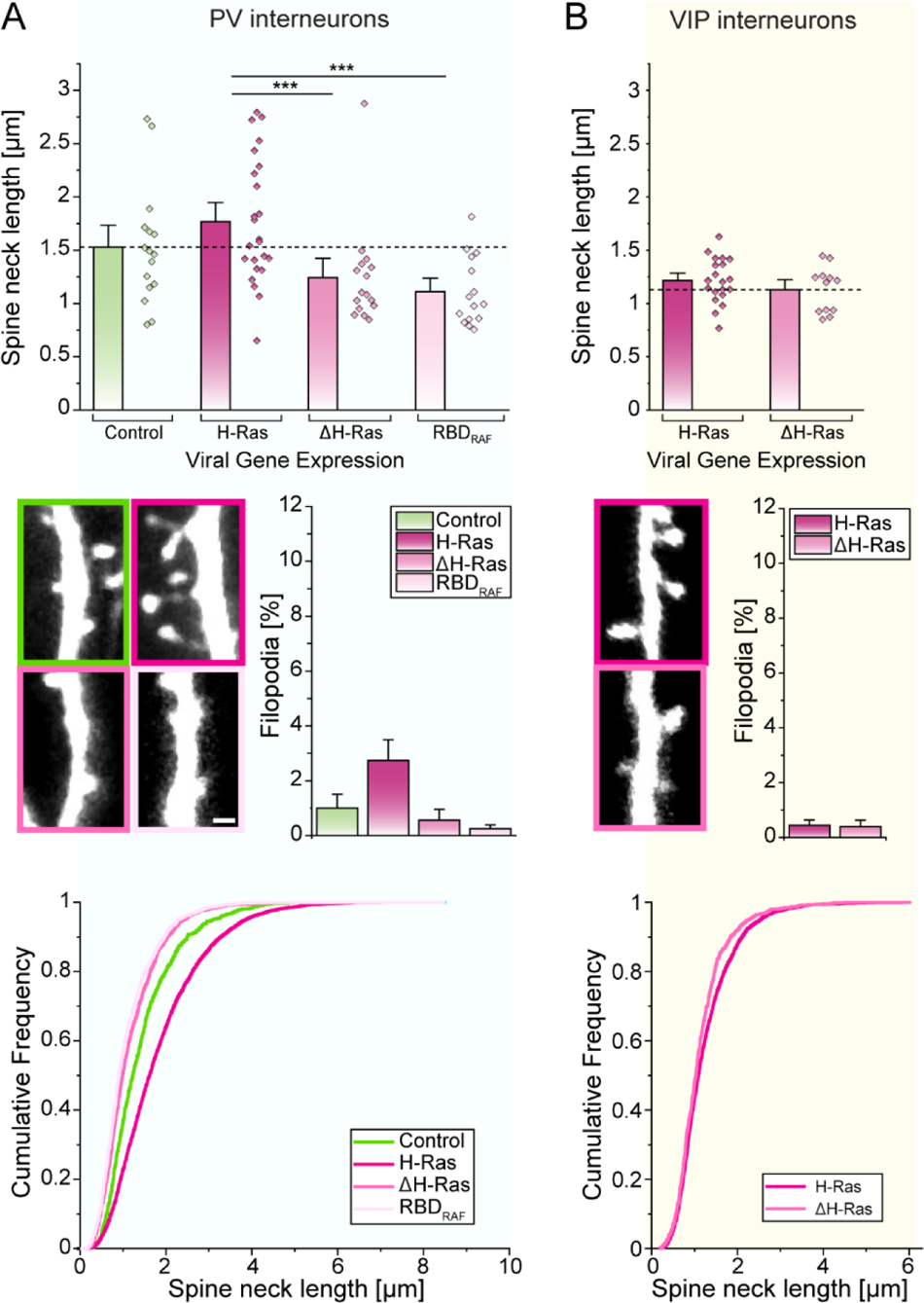
Increased spine neck length in H-Ras induced dendritic spines in interneurons. Visualization and analysis of spine morphology in PV-INs **(a)** and VIP-INs **(b)**. Control (green), H-Ras (magenta), ΔH-Ras (light magenta), and RBD_RAF_ (pink). (*Top*) A summary graph showing the spine neck length from individual neurons (dots) and the mean ± SEM spine neck length (bar). (*Middle*) Representative two-photon images of dendritic spines and their morphology. The percentage of filopodia at each expression condition. PV-INs expressing ectopic H-Ras have higher percentages of filopodia. (*Bottom*) Cumulative frequency plots showing the overall distribution of spine neck length. For PV-INs, Control: N= 16, H-Ras: N= 24, ΔH-Ras: N= 16, and RBD_RAF_: N= 15; For VIP-INs: H-Ras: N= 20, and ΔH-Ras: N= 12. ***P<0.001.

**Figure S5:**
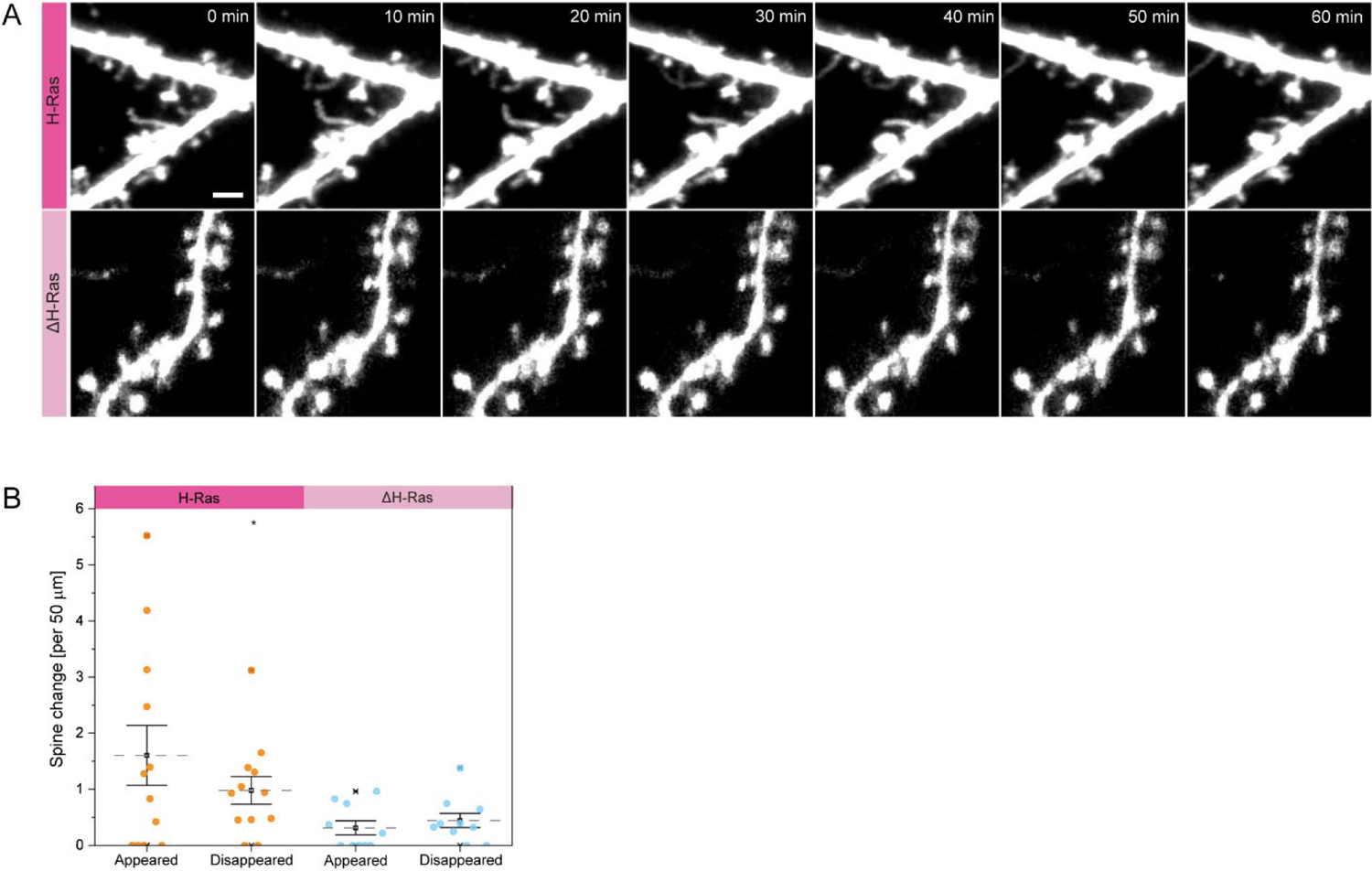
Increased spine dynamics in H-Ras induced dendritic spines. (a) Time-lapse two-photon images of H-Ras (top) and ΔH-Ras expressing neurons (bottom). Scale bar: 2 μm. (b) Analysis of spine dynamics (appearing, disappearing) in pyramidal neurons expressing ectopic H-Ras (magenta) and ΔH-Ras (light magenta). *P<0.05.

**Figure S6:**
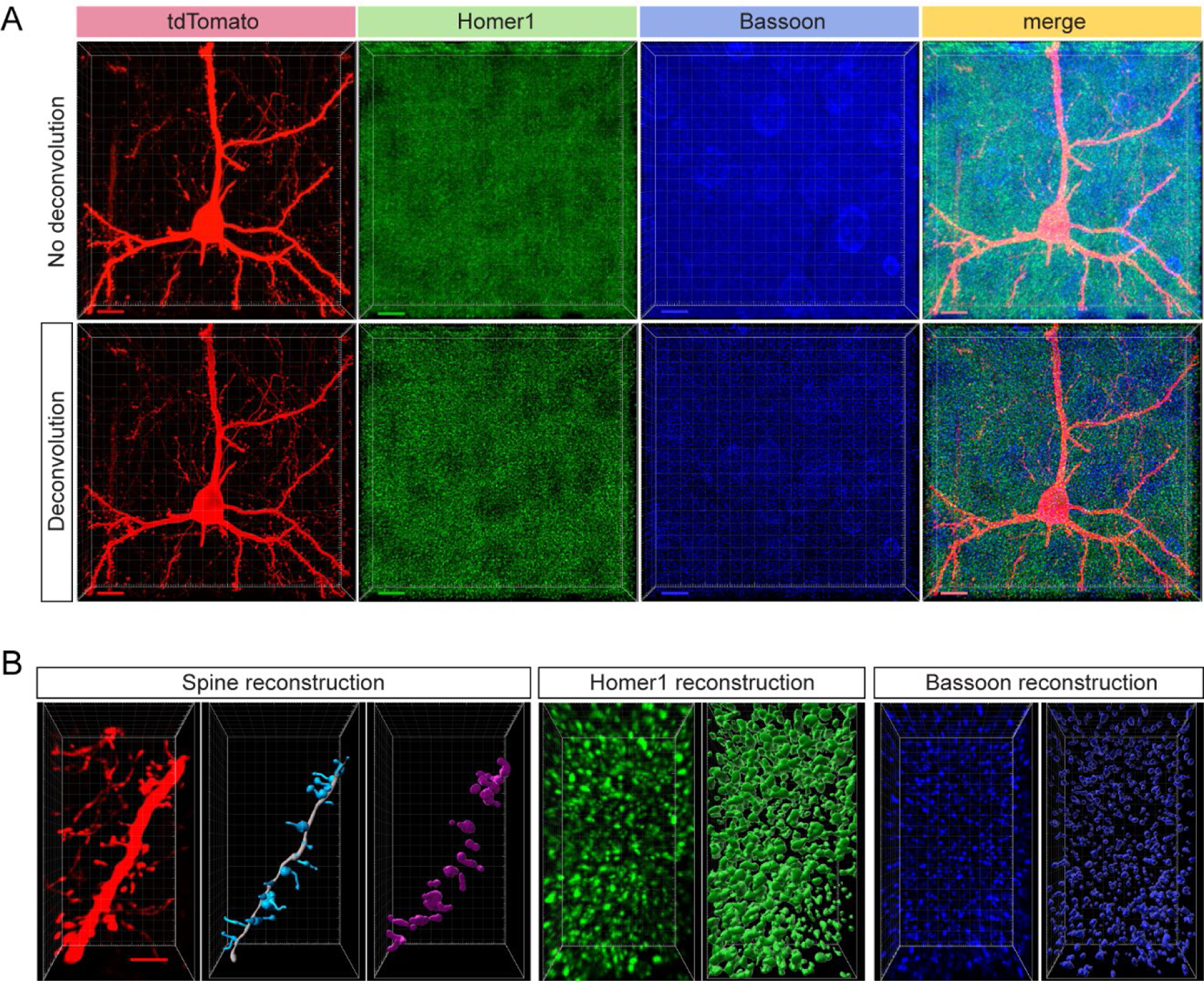
Deconvolution of antibody-stained confocal images. (a) Example of virus-injected and antibody-stained confocal images taken before and after deconvolution. Deconvolution improves signal-to-noise ratio in all channels (from left to right: tdTomato, Homer 1 stained against Alexa Fluor 633, bassoon stained against Alexa Fluor 405), merge). Scale bar 10 μm. (b) Representative reconstruction of spines, Homer1 and bassoon puncta via the Imaris software. Scale bar: 3 μm.

