## Supplementary Figures for "Exuberant *de novo* dendritic spine growth in mature neurons"

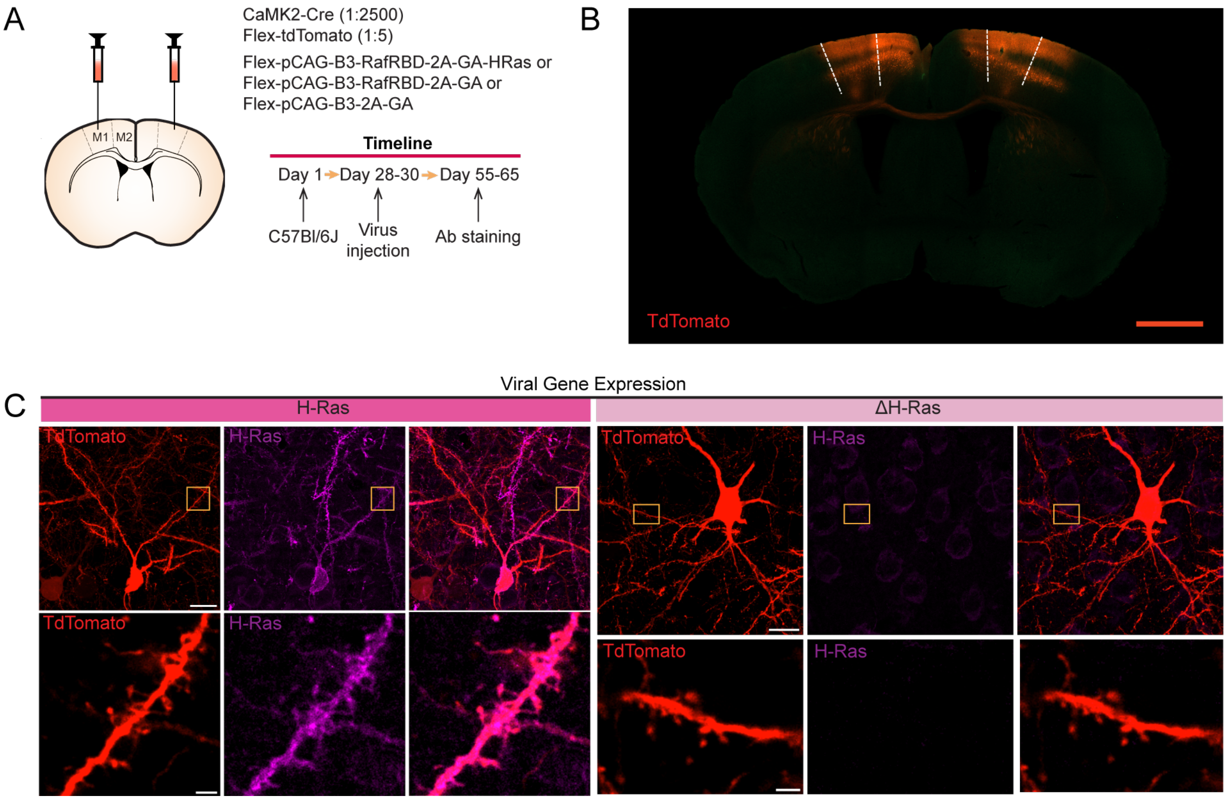


**Figure S1. Ectopic expression of H-Ras. (A)** Virus injection scheme and experimental timeline. **(B)** An example confocal overview image of tdTomato and ΔH-Ras expressing pyramidal cells at age ~P60 of a 40 μm fixed coronal brain slice. Scale bar: 1 mm. **(C)** Representative confocal image of pyramidal neurons (top) and magnified dendrites (bottom) of H-Ras (left) or a ΔH-Ras expressing cells (right). Images within the yellow box was zoomed in. From left to right: Neuron and dendrite images expressing the tdTomato, H-Ras protein stained with Alexa Fluor 633 (purple), and merge of channels. Scale bar: 10 μm (top), 2 μm (bottom).

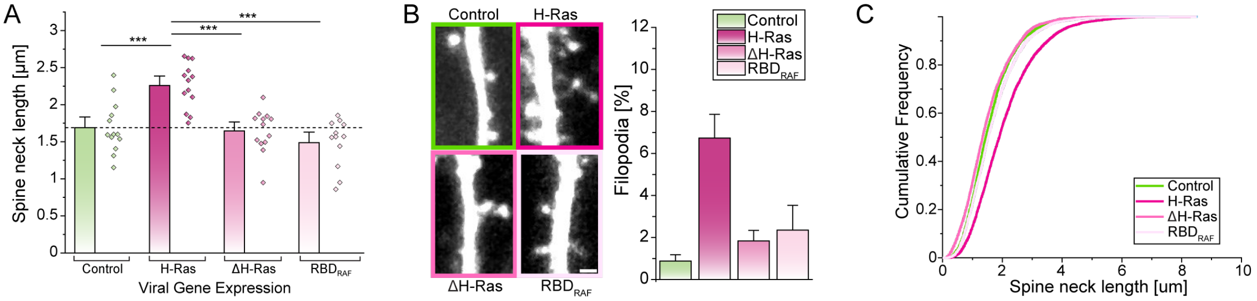


**Figure S2: Increased spine neck length in H-Ras induced dendritic spines. (A)** A summary graph showing the spine neck length of individual neurons (dots) and the mean ± SEM spine neck length (bar). Color scheme: Control (green), H-Ras (magenta), ΔH-Ras (light magenta), and $RBD_{RAF}$ (pink). **(B)** Representative two-photon images of dendritic spines and their morphology. The percentage of filopodia at each expression condition. Pyramidal neurons expressing ectopic H-Ras have higher percentages of filopodia. **(C)** A cumulative frequency plot showing the entire distribution of spine neck length. Control: N= 13, H-Ras: N= 13, ΔH-Ras: N= 15, and $RBD_{RAF}$: N= 10. ***P<0.001.

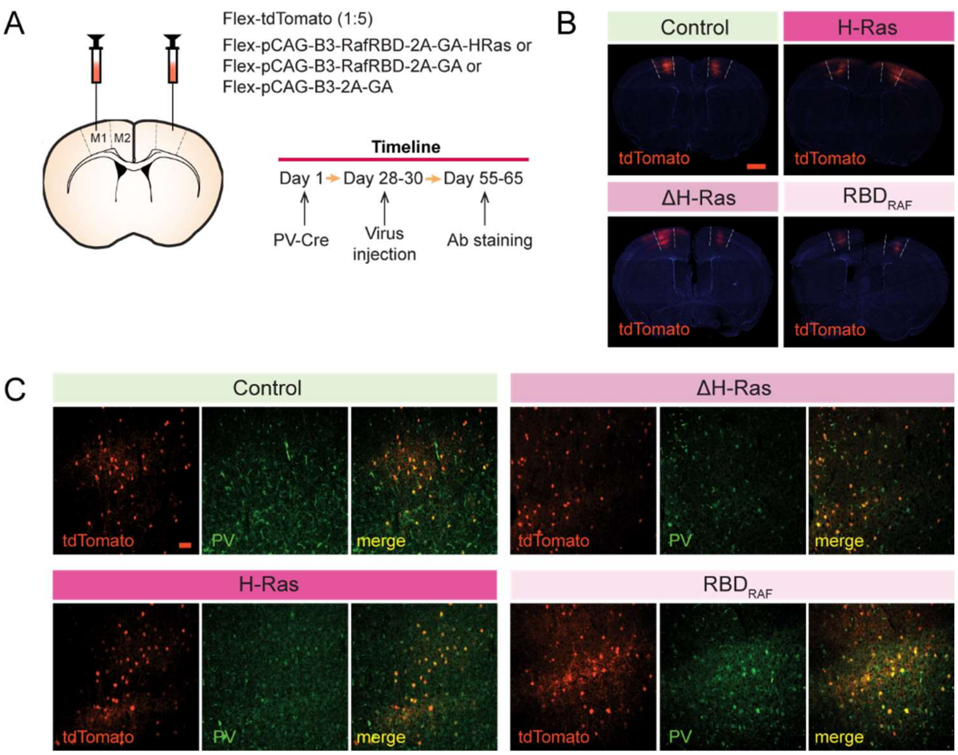


**Figure S3: Ectopic expression of H-Ras in PV-positive interneurons. (A)** Virus injection scheme and experimental timeline. **(B)** Example confocal images of pyramidal neurons expressing tdTomato (Control, top left) and either H-Ras (top right), ΔH-Ras (bottom left), or $RBD_{RAF}$ (bottom right) at age ~P60 of a 40 µm fixed coronal brain slice. Scale bar: 1 mm. **(C)** Representative confocal images showing that virus injected cells (red) are PV positive (green) stained with Alexa Fluor 633 no matter if they express tdTomato only (Control) (top left), ΔH-Ras (top right), H-Ras (bottom left), or $RBD_{RAF}$ (bottom right). Scale bar: 50 µm.

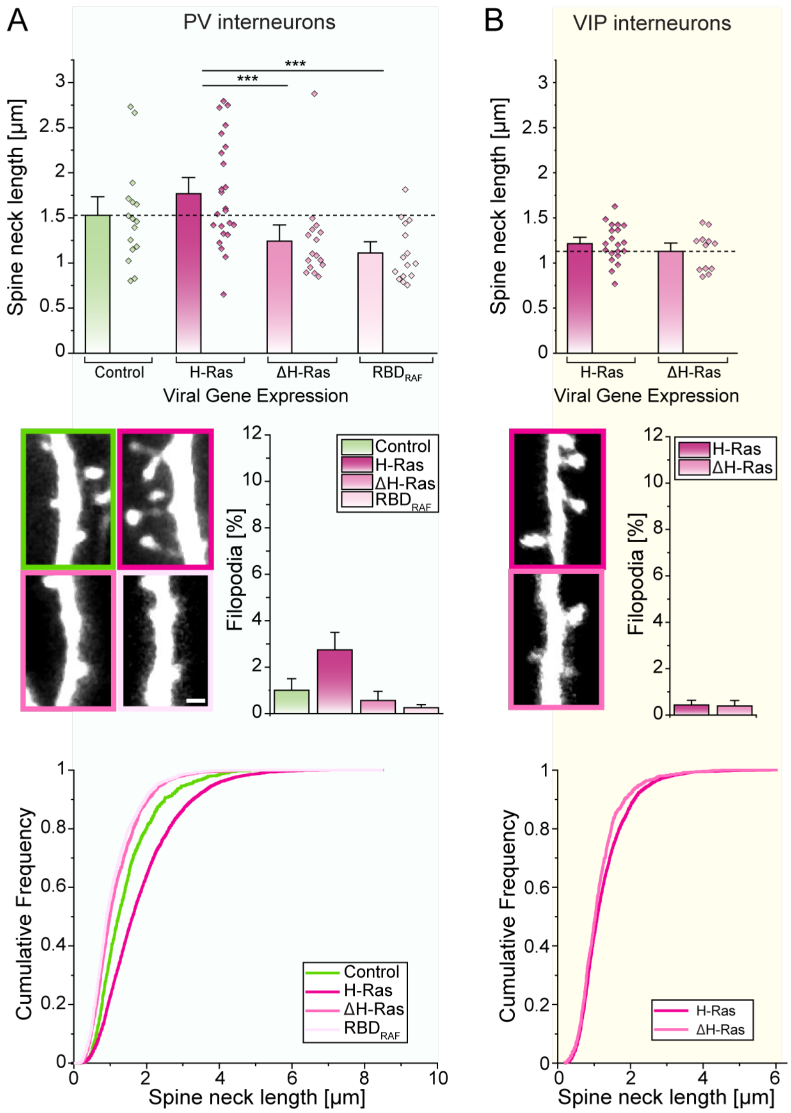


**Figure S4: Increased spine neck length in H-Ras induced dendritic spines in interneurons.** Visualization and analysis of spine morphology in PV-INs **(A)** and VIP-INs **(B)**. Control (green), H-Ras (magenta), ΔH-Ras (light magenta), and $RBD_{RAF}$ (pink). (Top) A summary graph showing the spine neck length from individual neurons (dots) and the mean ± SEM spine neck length (bar). (Middle) Representative two-photon images of dendritic spines and their morphology. The percentage of filopodia at each expression condition. PV-INs expressing ectopic H-Ras have higher percentages of filopodia. (Bottom) Cumulative frequency plots showing the overall distribution of spine neck length. For PV-INs, Control: N= 16, H-Ras: N= 24, ΔH-Ras: N= 16, and $RBD_{RAF}$: N= 15; For VIP-INs: H-Ras: N= 20, and ΔH-Ras: N= 12. ***P<0.001.

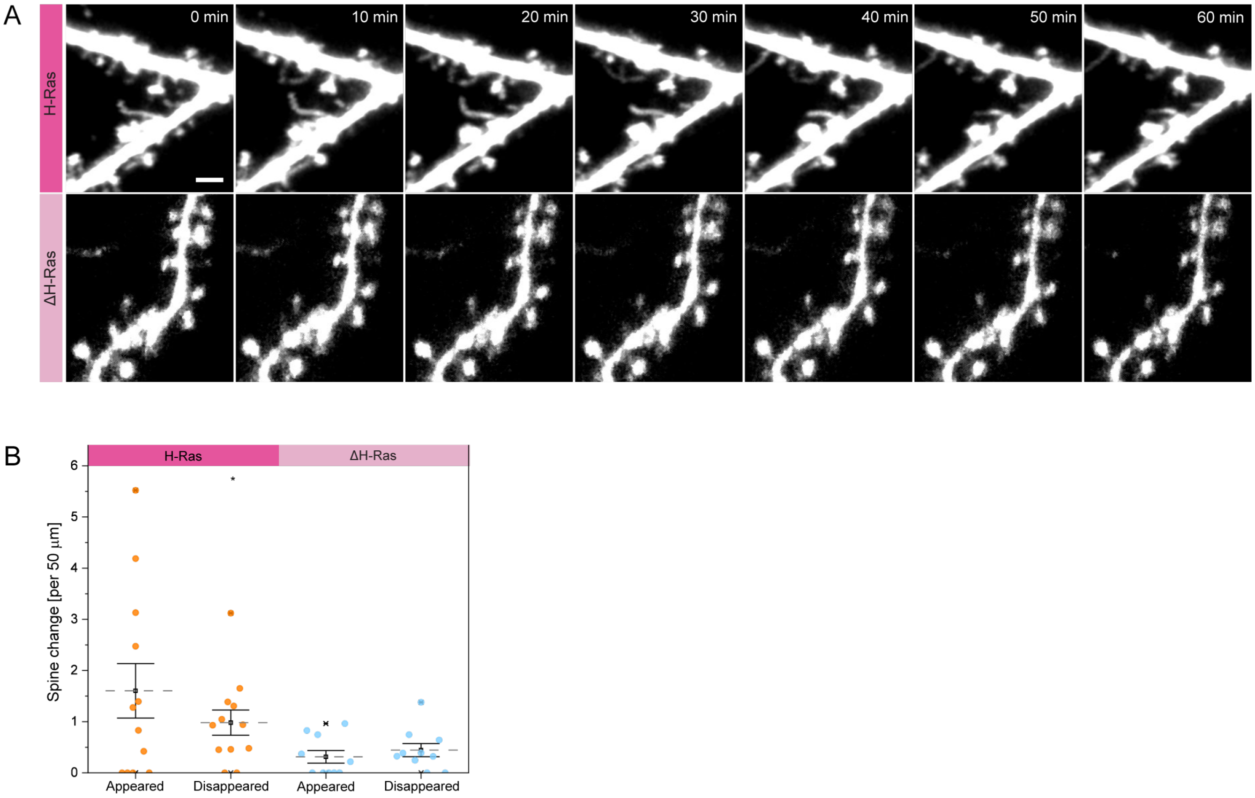


**Figure S5: Increased spine dynamics in H-Ras induced dendritic spines. (A)** Time-lapse two-photon images of H-Ras (top) and ΔH-Ras expressing neurons (bottom). Scale bar: 2 µm. **(B)** Analysis of spine dynamics (appearing, disappearing) in pyramidal neurons expressing ectopic H-Ras (magenta) and ΔH-Ras (light magenta). *P<0.05.

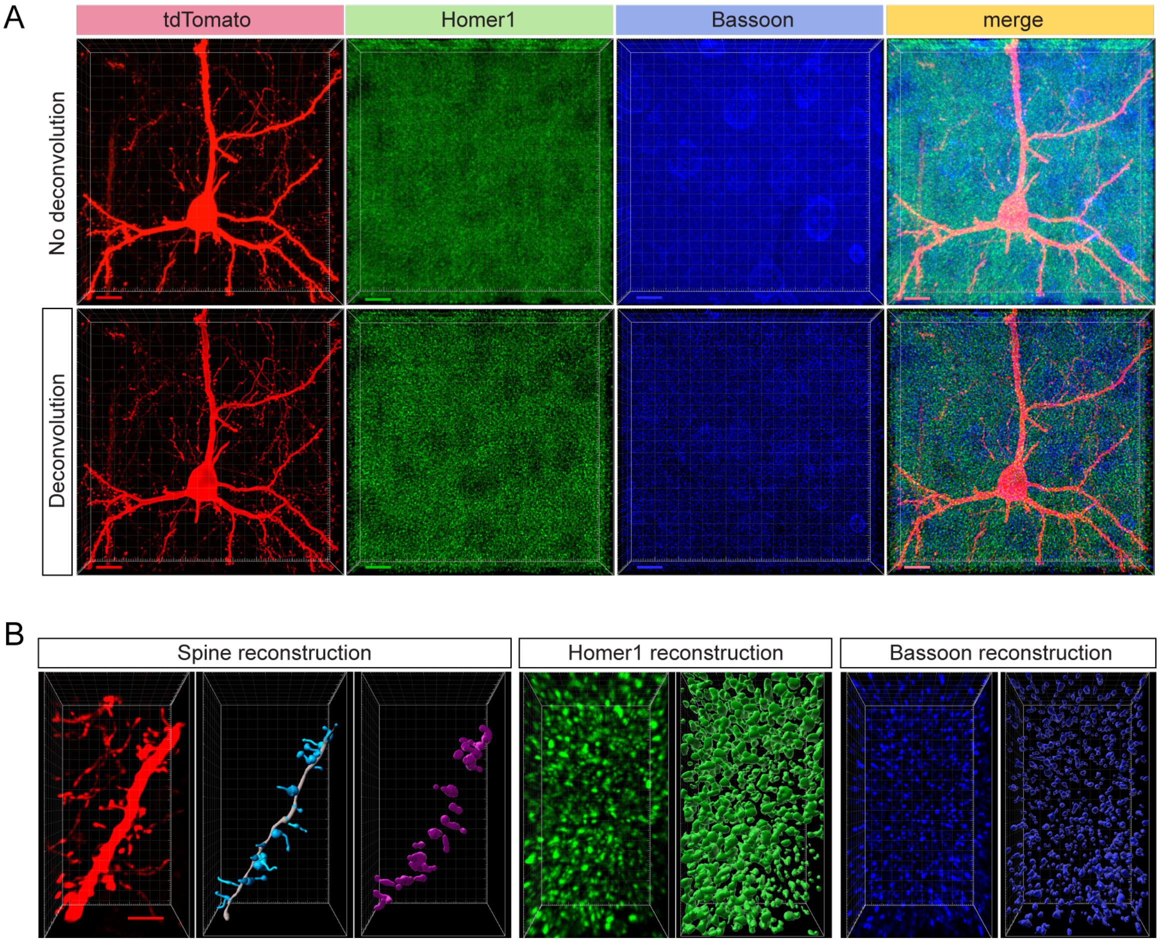


**Figure S6: Deconvolution of antibody-stained confocal images. (A)** Example of virus-injected and antibody-stained confocal images taken before and after deconvolution. Deconvolution improves signal-to-noise ratio in all channels (from left to right: tdTomato, Homer1 stained against Alexa Fluor 633, bassoon stained against Alexa Fluor 405), merge). Scale bar 10 μm. **(B)** Representative reconstruction of spines, Homer1 and bassoon puncta via the Imaris software. Scale bar: 3 μm.
